# AMPK regulates small RNA pathway prevalence to mediate soma-to-germ line communication and establish germline stem cell quiescence

**DOI:** 10.1101/2023.11.15.567172

**Authors:** Elena M. Jurczak, Christopher Wong, Shaolin Li, Fabian Braukmann, Ahilya N. Sawh, Thomas F. Duchaine, Eric A. Miska, Richard Roy

## Abstract

It is well established that cells communicate with each other via signaling molecules and pathways. Recent work has further indicated that this transfer of information can breach the soma-to-germ line barrier, thus permitting changes in germline gene expression in response to cellular decisions made in somatic lineages. We show that during periods of extended energy stress AMPK alters small RNA biogenesis in somatic cells, which non-autonomously regulates the quiescence of germline stem cells. By combining both genetic analyses and a novel method of miRNA imaging, we show that AMPK-mediated phosphorylation acts as a molecular switch that drives the re-allocation of the key RNA endonuclease Dicer to the miRNA synthesis pathway during the dauer stage of *C. elegans*. By modifying Dicer and other components of the miRNA synthesis machinery, AMPK fine-tunes the production of a population of somatic miRNAs that act as a “pro-quiescence” signal to maintain germline integrity during periods of extended energy stress, thus bridging the gap between the soma and the germ line by altering small RNA homeostasis.

## Introduction

Environmental fluctuations can impinge on cells to adopt context-specific gene expression programs, as to optimize energy homeostasis for growth, reproduction, or survival. This capacity of cells to adapt to their environment – termed “plasticity” – strongly contributes to reproductive fitness across organisms (Xiao et al., 2019). For example, when *C. elegans* larvae are exposed to various energetic stressors they can enter a phase of developmental quiescence (termed “dauer”), which allows them to preserve cellular integrity during intervals of suboptimal growth conditions (Hu, 2007). Following this challenge, the animals recover from the diapause with little to no consequence to their reproductive fitness. We have demonstrated that the highly conserved master regulator of metabolism AMP-activated protein kinase (AMPK) is required to coordinate the necessary molecular changes in both somatic tissues and germline stem cells to adapt to conditions of energetic stress (Narbonne & Roy, 2006; Narbonne & Roy, 2009). Upon activation, AMPK acts in somatic cells to influence the production of small RNAs that mediate a state of quiescence in germline stem cells during periods of stress (Kadekar & Roy, 2019; Wong et al., 2023). However, it remains unclear how AMPK interacts with small RNA pathways to mediate this effect, or how the regulation of small RNA production in the soma can promote the proper adaptation of germline stem cells to a given challenge.

The discovery of the first microRNA (miRNA) – *lin-4* – transformed our understanding of gene expression modulation by uncovering a class of regulators involved in the control of fundamental cellular and developmental processes (Lee et al., 1993; Wightman et al., 1993). In recent years, the vital relevance of miRNAs has emerged; the ability of individual miRNAs to suppress the expression of hundreds of transcripts in a tissue- and cell-specific manner confers a high level of plasticity to the cell (Xiao et al., 2019; O’Brien et al., 2018). Moreover, the dysregulation of miRNAs is prevalent in numerous diseases, and highly conserved miRNAs such as *let-7* represent important markers of cancer stem cells and tumourigenesis (Zhou et al., 2013).

It is now clear that gene expression changes in response to environmental conditions are mediated by a myriad of factors, including miRNAs and other small noncoding RNAs. Thousands of endogenous small noncoding RNAs that fall into three primary categories (small interfering RNAs (siRNAs), miRNAs, and Piwi-interacting RNAs (piRNAs)) enact the silencing of mRNA targets, thus enhancing the precision and speed of post-transcriptional regulation (Zhuang & Hunter., 2012; Billi et al., 2014). Indeed, one of the primary evolutionary advantages of small RNAs lies in their ability to respond rapidly to environmental stimuli and to reverse their mRNA target suppression upon return to basal conditions (Shimoni et al., 2007).

Despite the different functions of these classes of small noncoding RNAs, they share common factors required for their synthesis. The limited availability of these key factors may generate competition among the distinct biogenesis pathways (Loinger et al., 2012; Zhuang & Hunter, 2012). In *C. elegans*, for instance, DCR-1 (the orthologue of the endoribonuclease Dicer) is responsible for the synthesis of both miRNAs and of siRNAs; downstream, association of a small RNA with the appropriate Argonaute enables it to silence mRNA targets (Lee et al., 2006; Czech & Hannon, 2010; Zhuang & Hunter, 2012; Sawh & Duchaine, 2013). Through its association with different regulatory components, the endonuclease activity of DCR-1 can be directed to cleave pre-miRNAs and/or other double-stranded RNA substrates. These can then be further processed into the mature forms of miRNAs or siRNAs to execute their respective cellular tasks (Lee et al., 2013; Carthew & Sontheimer, 2009; Song & Rossi, 2017). The limited cellular abundance of DCR-1 may therefore contribute to competition among the pathways in certain physiological conditions, such that the active production of one class of DCR-1-dependent small RNAs proceeds at the expense of the production of the other classes (Zhuang & Hunter, 2012; Sawh & Duchaine, 2013).

To respond to shifts in resource availability, mechanisms have evolved in eukaryotic cells to optimize energy consumption and conservation. In particular, AMPK plays a critical role in integrating cues from the cellular environment. In response to conditions unfavourable for growth and development (such as lack of food or overcrowding), AMPK is activated by the kinase LKB1 (PAR-4 in *C. elegans*) and stimulates the production of ATP through the concerted activation of catabolic pathways and the inhibition of anabolic pathways (Herzig & Shaw, 2018). In this way, resources are conserved and ATP levels increase to meet the demands of cellular processes. Not surprisingly, the misregulation or absence of the LKB1/AMPK signalling axis leads to defects in metabolism, an inability to adapt to environmental stressors, and to impaired cell integrity and homeostasis (Narbonne & Roy, 2006; Narbonne & Roy, 2009).

In *C. elegans*, mutants that lack the AMPK catalytic subunits *aak-1* and *aak-2* (herein referred to as *aak(0)*) fail to halt germline stem cell proliferation upon entry into the dauer stage. Indeed, *aak(0)* dauer larvae either die or exhibit severe somatic and germline defects upon exit from this stage, including germline hyperplasia, sterility, and vulval defects (Narbonne & Roy, 2006; Narbonne & Roy, 2009; Kadekar & Roy, 2019). Compared to wild-type animals, AMPK mutants that transition through the dauer diapause display significantly altered epigenetic landscapes and modified germline gene expression profiles, indicating that these animals fail to adopt a dauer-specific gene expression program (Kadekar & Roy, 2019).

Recent findings indicate that AMPK affects chromatin modifications through its role in regulating the production of endogenous small RNAs in the soma (Kadekar & Roy, 2019). These small RNAs could potentially act across generations by contributing to the establishment of epigenetic modifications, such that the molecular memory specific to a given environmental condition is retained in subsequent generations (Duempelmann et al., 2020). Furthermore, the movement of small RNAs from the soma to the germ line indicate that these endogenous small RNAs might represent a preferred means of communication between these tissues (Conine & Rando, 2022).

In this study we describe a mechanism through which AMPK impinges upon DCR-1 to regulate the allocation of key enzymatic resources to different small RNA pathways, precisely altering small RNA homeostasis as cells adapt to conditions of energetic stress. We demonstrate that AMPK regulates the interaction between DCR-1 and the RNA-binding protein RBPL-1 to produce a population of miRNAs required for the establishment of germline stem cell quiescence during the dauer stage. This event in turn promotes germline integrity as animals transit through the dauer stage, ultimately preserving reproductive fitness. Our work thus reveals a regulatory mechanism that details how individual miRNAs can drastically alter the genetic and molecular profile of a critical population of stem cells and consequently control the reproductive fitness of the entire organism in response to environmental challenges.

## Results

### AMPK signalling regulates small RNA homeostasis as animals enter the dauer stage

Wild-type *C. elegans* larvae exposed to various stressors can survive in the developmentally quiescent dauer stage for months at a time with no negative consequences to their reproductive fitness. In contrast, *aak(0)* mutants either die during the dauer stage due to metabolic misregulation, or exhibit severe somatic and germline defects upon recovery, including germline hyperplasia and sterility (Narbonne and Roy 2006; Narbonne and Roy 2009; Kadekar & Roy, 2019). The compromise of small RNA biogenesis pathway components RDE-4 and ERGO-1 partially suppresses the germline defects in AMPK mutants (Kadekar & Roy, 2019); we thus hypothesized that AMPK likely impinges on factors involved in the synthesis of small RNAs in the soma to send a pro-quiescence signal to germline stem cells at the onset of the dauer stage.

To test this hypothesis, we performed small RNA sequencing on *daf-2* and *daf-2; aak(0)* dauer larvae (herein referred to as “control” and *“aak(0)”*, respectively) to determine whether the expression profile of endogenous small RNAs is altered in AMPK mutants that transit through the dauer stage. This analysis identified dramatic and global changes in the expression of endogenous small RNAs. Most interestingly, we found that the ALG-3/4 class 26G siRNAs are largely upregulated in AMPK dauer mutants, while the miRNAs are inversely downregulated (Fig. 1A; Fig. S1A-C). These contrasting changes in expression led us to wonder whether AMPK may be involved in allocating enzymatic resources to the miRNA and siRNA biogenesis pathways, as to favour the production of one class of small RNAs over another as an adaptive response to a specific environmental cue.

**Figure 1:**
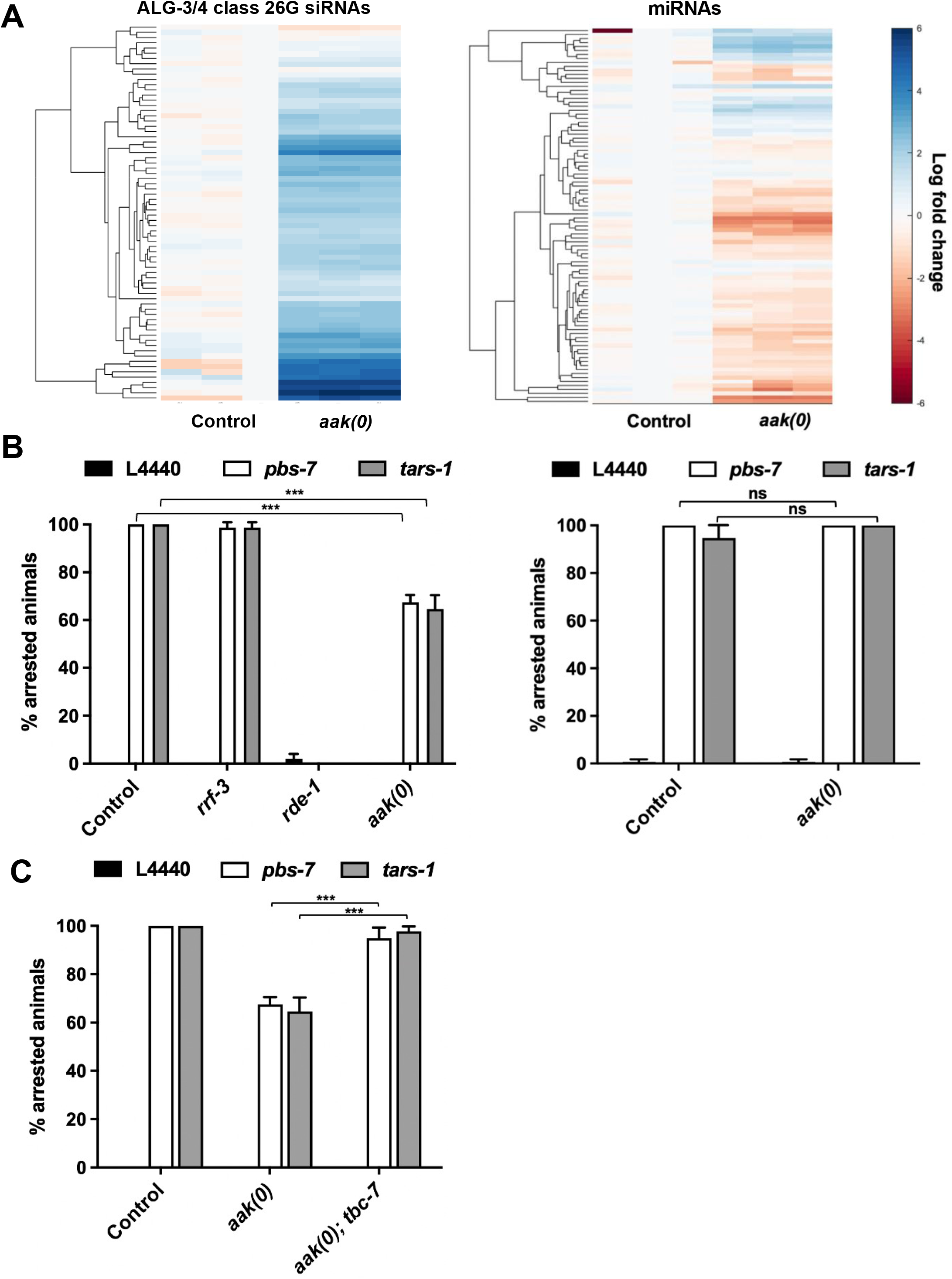
Loss of AMPK signalling results in the dramatic perturbation of small RNA homeostasis. (A) Heat map representation of small RNA sequencing analysis demonstrates the relative upregulation of 26G-siRNAs and the global downregulation of miRNAs in *daf-2; aak(0)* dauer mutants compared to *daf-2* controls. Clustering was performed using the Matlab function “clustergram” and calculates euclidian distance. (B-C) Animals were fed with arrest-inducing dsRNA targeting the genes *pbs-7* and *tars-1*. B) Animals were then allowed to transit through and recover from the dauer stage (left) or to grow at the permissive temperature without developmental interruptions (right), and the percentage of arrested animals was quantified. (C) Animals were allowing to transit through and recover from the dauer stage, and the percentage of post-dauer arrested animals was quantified. *\*\*\*P* < 0.001 and *\*P* < 0.05, when compared to L4440 (RNAi empty vector) control using two-way ANOVA and Holm-Šidák correction. Data are represented as mean ± SD. All animals carry the *daf-2(e1370)* allele. n ≥ 50. See also Figure S1.

To determine whether AMPK dauer larvae exhibit deregulated competition between small RNA pathways, an RNAi sensitivity assay was designed to assess the capacity of these animals to carry out exogenous RNAi. Animals were fed dsRNA targeting *pbs-7* (*<u>p</u>roteasome <u>b</u>eta <u>s</u>ubunit-7*) and *tars-1* (*<u>tR</u>NA <u>s</u>ynthetase-1*) – two genes whose compromise induces early larval arrest in 100% of control animals transiting through the dauer stage. In contrast, when AMPK mutants are subjected to these dsRNA and dauer entry is induced, a sizeable portion of the animals bypass larval arrest and reach adulthood upon recovery (Fig. 1B). AMPK mutants which do not transit through the dauer stage, however, exhibit full RNAi sensitivity (Fig. 1B). Altogether, these results indicate that AMPK mutants placed in conditions of stress which induce the dauer diapause are partially <u>R</u>NAi <u>de</u>fective (Rde).

We recently showed that the compromise of the RabGAP *tbc-7* suppresses the germline hyperplasia and post-dauer sterility of AMPK mutants (Wong et al., 2023). To determine if the relationship between the partial <u>Rde phenotype</u> and the post-dauer sterility observed in AMPK mutants are linked, we compared the RNAi sensitivity of *aak(0)* animals with that of *aak(0); tbc-7* mutants that exhibit a restoration of post-dauer fertility. The RNAi sensitivity is restored to wild-type-like levels in the *aak(0); tbc-7* mutants, suggesting that suppressing the germline defects associated with a lack of AMPK signalling correlates with restoring RNA homeostasis (Fig. 1C).

These results suggest that AMPK is required to establish a specific endogenous small RNA expression profile upon exposure to energetic stress. In the absence of AMPK signalling, endogenous siRNAs (endo-siRNAs) are upregulated, miRNAs are downregulated, and animals fail to appropriately process exogenous sources of dsRNA, rendering the animals partially RNAi defective.

### DCR-1 abundance is limiting in animals that lack AMPK

We surmised that the inappropriate regulation of Dicer (DCR-1), which is required for the synthesis of many endogenous and exogenous small non-coding RNAs, might account for the observed disruption in small RNA homeostasis in the AMPK mutants. If the activity of DCR-1 is simultaneously solicited by multiple small RNA synthesis pathways, it could become limiting, potentially impacting the outcomes of one or more small RNA-dependent processes. Therefore, at the onset of the dauer stage, AMPK may be required to re-allocate DCR-1 activity to the pathway(s) where its contribution is most urgent.

If this is indeed the case, then increasing the availability of DCR-1 should reduce or eliminate the competition for this enzyme among the various biosynthesis pathways, permitting the production of all classes of DCR-1-dependent small RNAs. Consistent with this, driving the expression of *dcr-1* ubiquitously in the soma of AMPK mutants led to a significant suppression of post-dauer sterility (Fig. 2A). The somatic defects typically observed in these animals remained elevated, likely due to the misexpression of small RNAs involved in development, and suggesting that DCR-1 acts cell non-autonomously in the soma to specifically control germline integrity (Fig. 2B).

**Figure 2:**
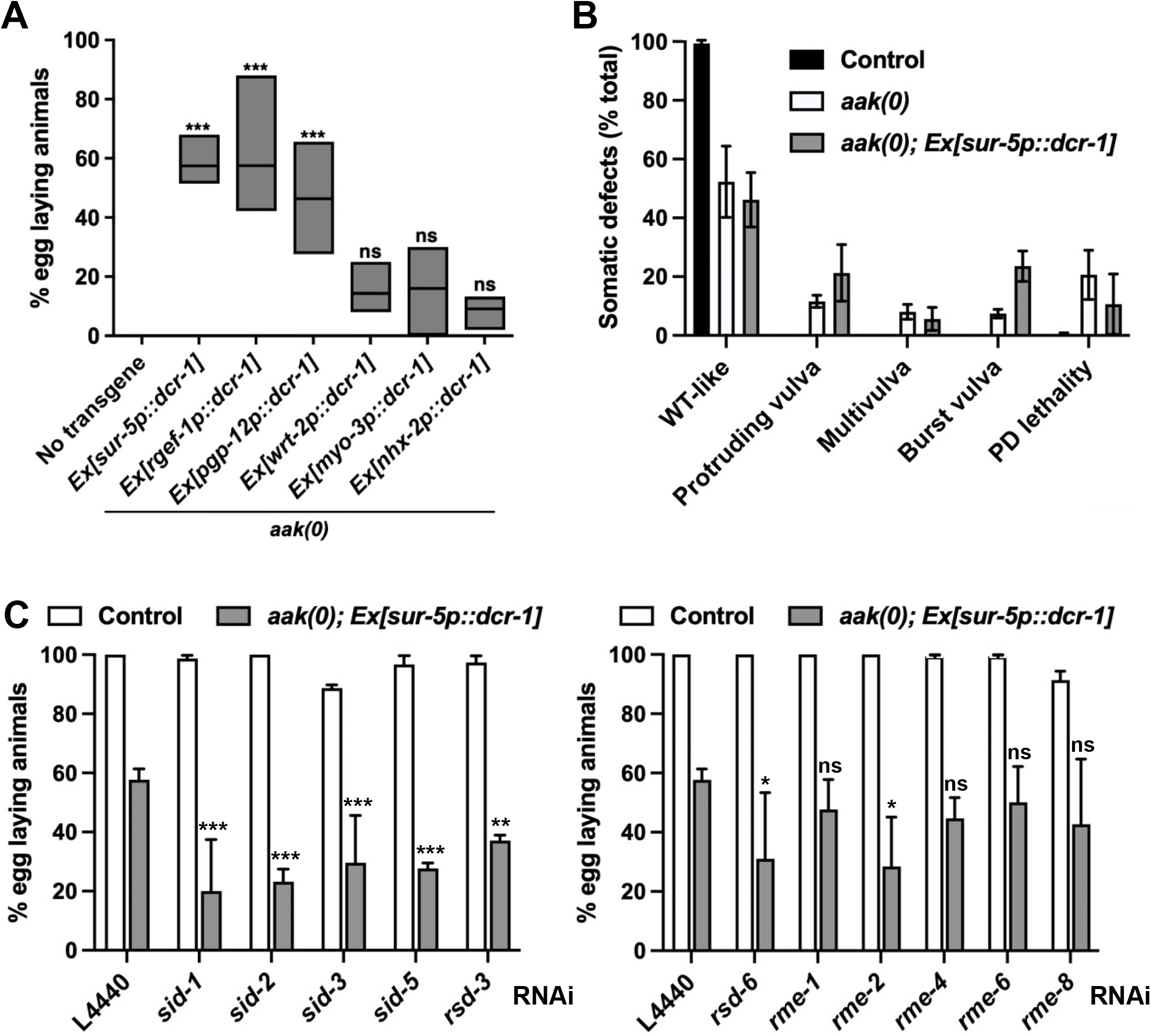
DCR-1 activity is limiting in AMPK mutants. (A) The expression of *dcr-1* was driven in *aak(0)* mutants using the somatic tissue-specific promoters *sur-5p* (all somatic cell types)*, rgef-1p* (nervous system)*, pgp-12p* (excretory system), *wrt-2p* (hypodermis)*, myo-3p* (body wall muscle), and *nhx-2p* (intestine), and post-dauer fertility was assessed. *\*\*\*P* < 0.001, *\*\*P* < 0.01, and *\*P* < 0.05 when compared to *aak(0)* control using one-way ANOVA and Holm-Šidák correction. (B) Assessment of the post-dauer somatic defects observed in AMPK mutants that express a *dcr-1* transgene (*aak(0); Ex[sur-5p::dcr-1])*. (C) The expression of members of the *sid* family (dsRNA transport), *rme* family (endocytosis), and *rsd* family (dsRNA spreading) were compromised using RNAi in AMPK mutants that express a *dcr-1* transgene, and post-dauer fertility was assessed. *\*\*\*P* < 0.001, *\*\*P* < 0.01, and *\*P* < 0.05 when compared to L4440 control using two-way ANOVA and Holm-Šidák correction. Data are represented as mean ± SD. All animals carry the *daf-2(e1370)* allele. n ≥ 50.

AMPK has been shown to act in the neurons and in the excretory system to establish quiescence (Kadekar & Roy, 2019). Therefore, we wondered whether AMPK may target DCR-1 in these tissues to establish a small RNA profile consistent with the changes in gene expression that must occur in animals as they enter the dauer stage. We provided additional copies of this enzyme by expressing *dcr-1* in various somatic tissues to determine if it might be required in the same tissues as AMPK. In addition, if increasing the abundance of *dcr-1* in these tissues is sufficient to correct the AMPK mutant germline deficiencies, we could conclude that its activity is limiting in these animals, and that AMPK works directly, or indirectly, through *dcr-1* to control the viability of germ cells during the dauer stage.

A modest improvement in post-dauer fertility was observed by expressing *dcr-1* from multicopy transgenes in the hypodermis, in the body wall muscle, and in the intestine of transgenic animals. In contrast, driving the expression of *dcr-1* in the nervous system and in the excretory system led to strong rescues of post-dauer sterility, indicating that AMPK and DCR-1 most likely act in the same somatic tissues to establish germline stem cell quiescence as animals enter the dauer stage (Fig. 2A).

To enhance our understanding of how *dcr-1* could regulate the germline non-autonomously, we explored the requirement for the downstream transport of small RNAs produced by DCR-1. The movement of different types of RNA between cells is mediated through designated dsRNA importers and exporters. In *C. elegans*, systemic RNAi and the transport of dsRNA rely on the *sid* (<u>S</u>ystemic RNAi <u>D</u>efective)*, rsd* (<u>R</u>NAi <u>S</u>preading <u>D</u>efective), and *rme* (<u>R</u>eceptor-<u>M</u>ediated <u>E</u>ndocytosis) families of genes (Saint-Pol et al., 2004; José, 2015; Marré et al., 2016; Wang & Hunter, 2017). We questioned whether these factors may similarly be involved in the transport of double stranded RNA produced downstream of DCR-1 processing at the onset of the dauer stage. Using RNAi we disabled the expression of the best characterized *sid, rsd,* and *rme* genes in *aak(0)* mutants that express additional copies of DCR-1. Though the loss of the *rme* genes had little to no effect on the *dcr-1*-dependent post-dauer fertility in these animals, the compromise of the *sid* genes and of *rsd-3* reduced fertility, despite the increased abundance of *dcr-1* (Fig. 2C). Altogether, these data indicate that the suppression of AMPK post-dauer germline defects conferred by the overexpression of *dcr-1* is dependent upon the proper transport and spreading of dsRNA between and/or within tissues.

### Small RNA biogenesis pathways compete for Dicer during the dauer stage

Each class of small RNA requires a devoted set of factors for its production, while the loss of one or more of these key components often results in defective synthesis and the corresponding physiological and/or developmental phenotypes in the animal. To determine if a specific class of endogenous small RNA is required for establishing germline integrity during dauer, we used RNAi to compromise the expression of key components unique to each of the pathways involved in small RNA biogenesis. We reasoned that by inhibiting the production of small RNAs unrelated to the establishment of quiescence, we could consequently increase the availability of limiting components, namely DCR-1, by reducing competition for its activity. As a result, DCR-1 function could be re-allocated toward the synthesis of specific classes of pro-quiescent small RNAs, thereby restoring germline integrity.

To test this hypothesis, we explored the roles of key required components of the endo-siRNA biogenesis machinery: the DCR-1 interactor RDE-4 and the ERI complex, (Duchaine et al., 2006; Billi et al., 2014). Compromising the expression of these factors led to the partial suppression of AMPK post-dauer sterility (Fig. 3A), indicating that the endo-siRNA biogenesis axis must normally be attenuated as animals prepare to enter the dauer stage. Conversely, any gain in fertility conferred by disabling endo-siRNA biosynthesis was diminished following RNAi against *drsh-1* or *pash-1*, members of the Microprocessor complex required for miRNA biogenesis (Denli et al., 2004) (Fig. 3B). These results suggest that miRNAs are required to protect germ cell integrity during the diapause.

**Figure 3:**
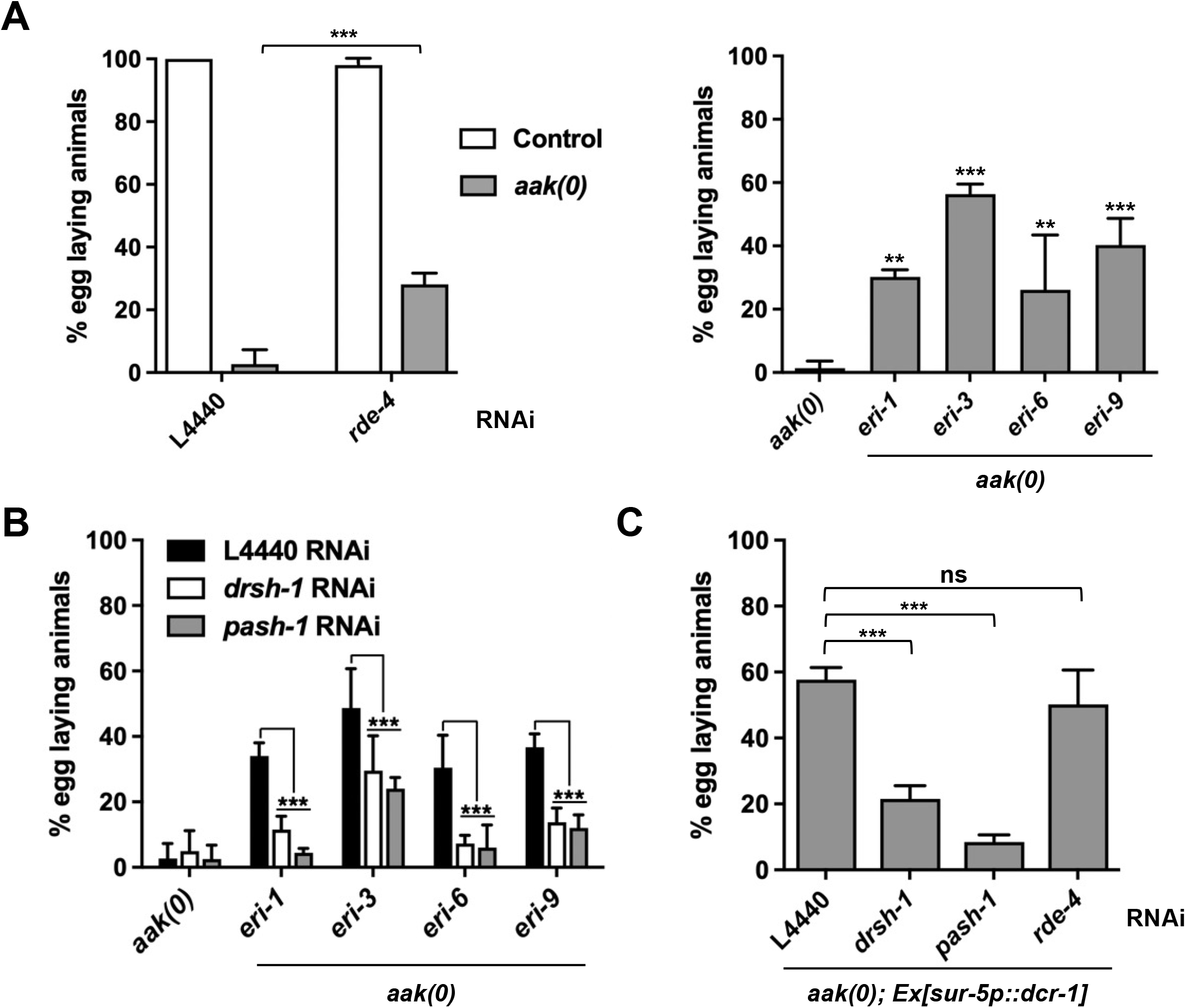
Small RNA biogenesis pathways compete for limited DCR-1 availability. (A) The expression of factors required for siRNA biogenesis was compromised using RNAi (left) or genetic deletions (right), and post-dauer fertility was assessed. (B) *eri* mutants were subjected to *drsh-1* and *pash-1* RNAi, and post-dauer fertility was assessed. (C) RNAi against *drsh-1*, *pash-1*, or *rde-4* was performed in AMPK mutants expressing a *dcr-1* transgene (*aak(0); Ex[sur-5p::dcr-1])*, and post-dauer fertility was assessed\*\*\**P* < 0.001, *\*\*P* < 0.01, *\*P* < 0.05 when compared to control using one-way ANOVA or two-way ANOVA and Holm-Šidák correction. Data are represented as mean ± SD. All animals carry the *daf-2(e1370)* allele. n ≥ 50.

Furthermore, the compromise of miRNA biogenesis in the *dcr-1*-expressing AMPK mutants reduced the fertility of the animals substantially, while disrupting endo-siRNA production in the same animals had no such effect (Fig. 3C). Taken altogether, these results are consistent with an important role for miRNA biogenesis in the protection and preservation of the germline stem cells during lengthy periods of starvation. Moreover, the production of pro-quiescent miRNAs appears to be favoured by the AMPK-dependent re-allocation of DCR-1 away from the production of further categories of small RNAs.

### AMPK drives Dicer re-allocation and defines pathway selection at the onset of the dauer stage

Our findings prompted us to explore whether the AMPK-mediated regulation of one or more factors involved in siRNA and miRNA biogenesis could toggle DCR-1 to redirect its activity from one pathway to another. This could potentially be achieved by altering the association between DCR-1 and either of its two primary interactors: the dsRNA-binding proteins RBPL-1 and RDE-4 (*C. elegans* homologs of the mammalian PACT/ RBBP6 and TRBP, respectively). Previous work has shown Dicer to preferentially associate with TRBP, thus favouring siRNA biogenesis by default (Lee et al., 2013). In contrast, the complex formed by Dicer and PACT/RBBP6 leads to the predominant production of miRNAs (Lee et al., 2013).

In light of our results with increased *dcr-1* abundance, we reasoned that if we could tip the balance in the competition for DCR-1 function by expressing additional copies of *rbpl-1* ubiquitously in the soma of *aak(0)* mutants, we could generate a supraphysiological abundance of RBPL-1 compared to RDE-4, favouring the association between DCR-1 and RBPL-1. Consistent with this scenario, the ubiquitous somatic expression of RBPL-1 suppressed post-dauer sterility significantly, leading us to infer that the production of miRNAs is favoured through the RBPL-1-mediated sequestration of DCR-1 for miRNA biogenesis (Fig. 4A). Both *drsh-1* and *pash-1* RNAi revert the restored fertility, consistent with a requirement for RBPL-1 and miRNA production in the preservation of reproductive fitness (Fig. 4A).

**Figure 4:**
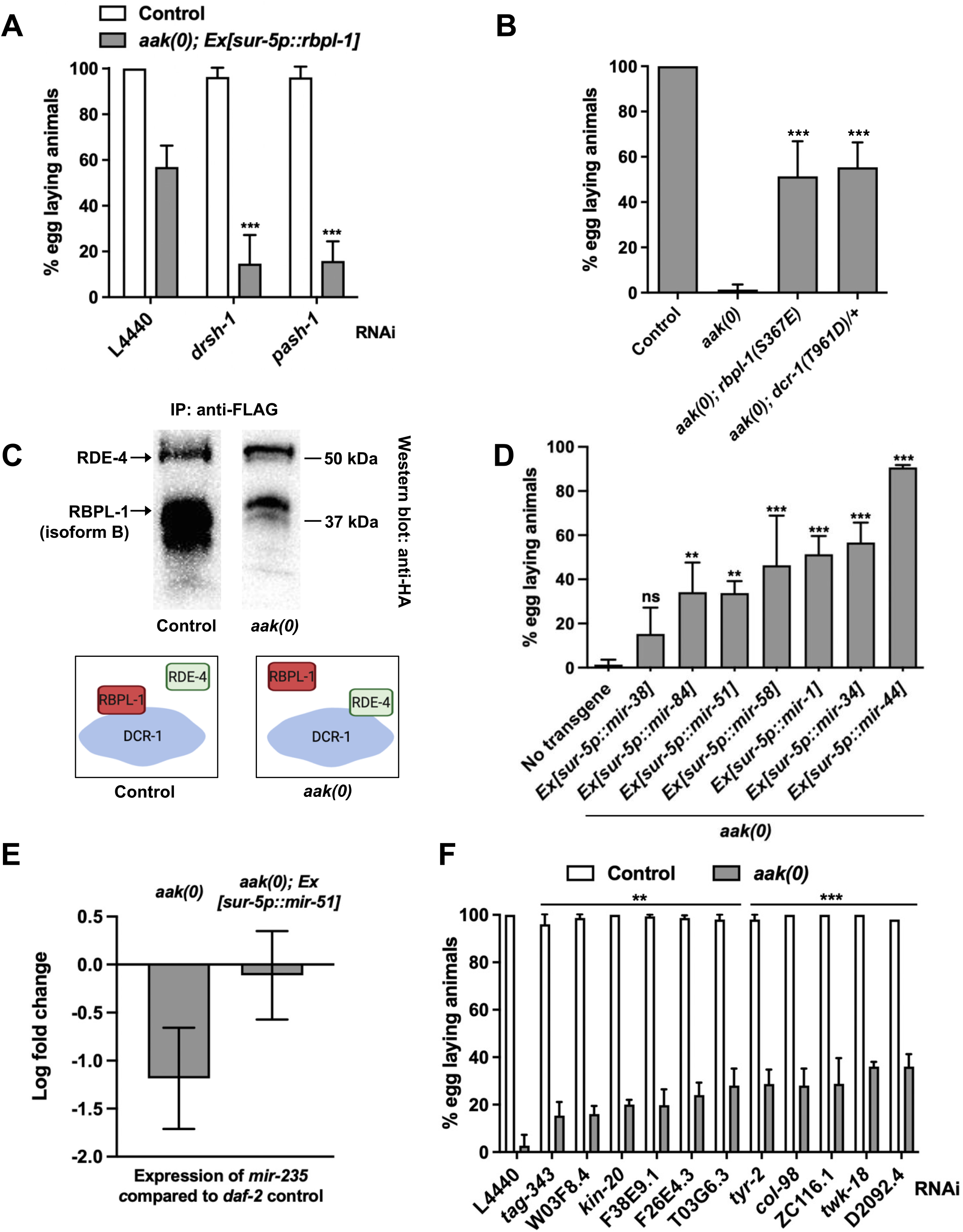
AMPK targets DCR-1 and RBPL-1 to promote miRNA production and maintain post-dauer fertility. (A) The expression of *rbpl-1* was driven ubiquitously in the somatic tissues of *aak(0)* animals using a *sur-5p* promoter. Animals were subjected to *drsh-1* and *pash-1* RNAi, and post-dauer fertility was assessed. \*\*\**P* < 0.001, when compared to L4440 control using two-way ANOVA and Holm-Šidák correction. (B) The phosphomimetic mutations RBPL-1(S367E) and DCR-1(T961D)/+ were generated in *aak(0)* mutants using CRISPR-Cas9 genome editing, and post-dauer fertility was assessed. *\*\*\*P* < 0.001 when compared to *aak(0)* control using one-way ANOVA and Holm-Šidák correction. (C) Top: Immunoprecipitation using an anti-FLAG antibody was performed on whole protein extracts from wild-type (left) and *aak(0)* (right) dauer larvae that express DCR-1::FLAG (tagged endogenously using CRISPR-Cas9). RBPL-1 and RDE-4 were tagged with HA using CRISPR-Cas9 and expressed endogenously in both genetic backgrounds. Membranes were probed with antibodies that recognize the HA epitope. Bottom: graphical summary of co-immunoprecipitation experiment demonstrating changing DCR-1 interaction dynamics in the presence and absence of AMPK signalling. (D) Post-dauer fertility was assessed after the indicated miRNAs were expressed ubiquitously using a somatic promoter (*sur-5p*) in *aak(0)* mutants. *\*\*\*P* < 0.001 and *\*P* < 0.05 when compared to *aak(0)* control using one-way ANOVA and Holm-Šidák correction. (E) Changes in the expression levels of *mir-235* were measured in control, *aak(0)*, and *aak(0); Ex[sur-5p::mir-51]* dauer larvae using RT-qPCR analysis. (F) An RNAi survey was performed against predicted mRNA targets of *mir-34* in *aak(0)* mutants and post-dauer fertility was assessed. *\*\*\*P* < 0.001 and *\*\*P* < 0.01 when compared to L4440 control using two-way ANOVA and Holm-Šidák correction. Data are represented as mean ± SD. All animals carry the *daf-2(e1370)* allele. n ≥ 50. See also Figures S2-3 and Tables S1-3.

The activity of AMPK in the cell is typically mediated through the sequence-specific phosphorylation of its targets. We therefore examined the primary sequences of DCR-1 and of all DCR-1-dependent, small RNA biogenesis factors, including RBPL-1, for the presence of AMPK phosphorylation motifs. From our bioinformatic analysis, two candidates stood out, each of which possessed putative high stringency AMPK phosphorylation motifs: DCR-1 and RBPL-1 (Table S1). We generated phosphomimetic variants of both proteins using CRISPR-Cas9 genome editing, and found that these mutations (DCR-1(T961D)/+ and RBPL-1(S367E)) led to the significant restoration of post-dauer fertility in *aak(0)* mutants (Fig. 4B; Fig. S2A-B), seemingly suggesting that AMPK acts on DCR-1 and RBPL-1 to control small RNA biogenesis during the dauer stage.

To gain understanding of this mechanism, DCR-1 (DCR-1::FLAG) was immunoprecipitated in control and *aak(0)* dauer larvae, and the recovered fractions were probed to detect RBPL-1 (RBPL-1::HA) and RDE-4 (RDE-4::HA). The levels of RDE-4 were comparable in both genetic backgrounds, but although the long isoform of RBPL-1 was not detected in either IP sample, we detected high levels of the short isoform of RBPL-1 (whose predicted molecular weight is of 42.6 kDa) in the *daf-2* control background. This isoform was also present in the AMPK mutant background, albeit at significantly reduced levels (Fig. 4C; Fig. S3A-B), consistent with our hypothesis which postulated that AMPK favours an interaction between DCR-1 and RBPL-1. Taken together, these results suggest that AMPK re-configures the interactions between Dicer and its functional interacting partners involved in small RNA biogenesis by triggering the timely re-allocation of this critical endonuclease to the miRNA biogenesis pathway through its association with RBPL-1.

### A subset of somatic miRNAs produced downstream of AMPK promote germline integrity during the dauer stage

Small RNA-sequencing and genetic analysis both suggest that the inability of germline stem cells to establish quiescence during the dauer stage arises from global changes in miRNA homeostasis in the AMPK mutants. It is unclear however, whether individual miRNAs or entire miRNA families may be specifically required to target germ cell transcripts to maintain germline integrity as animals adapt to metabolic stress and arrest development.

Categorizing the miRNAs in our dataset according to their function and degree of downregulation led to the identification of a subset of candidates which could potentially mediate the cell cycle arrest and quiescence programs typical of the dauer diapause (Alvarez-Saavedra & Horvitz, 2010) (Table S2). To test if any of these miRNAs could correct the germline integrity of AMPK mutant dauer larvae, the expression levels of individual miRNAs that belong to several of these families were restored in the soma of AMPK mutants. This allowed us to identify pre-miRNAs, the transgenic expression of which significantly suppressed the post-dauer sterility of *aak(0)* mutants (Fig. 4D); these results suggest that the expression of miRNAs in somatic cells is sufficient for transmitting a pro-quiescence signal to germline stem cells.

We wished to determine whether this phenotype resulted from the individual requirement of the specific miRNAs tested, or whether the increased abundance of pre-miRNA substrate could favour miRNA biogenesis through Dicer re-allocation. The expression levels of an arbitrary miRNA (*mir-235*) were measured in control, *aak(0)*, *and aak(0); Ex[sur-5p::mir-51]* dauer larvae using RT-qPCR. As previously shown through small RNA sequencing analysis, the levels of *mir-235* were downregulated in *aak(0)* mutants. In contrast, they returned to near wild-type levels in mutants expressing additional copies of *mir-51* (Fig. 4E). These results confirm that the defects caused by loss of AMPK signalling can be corrected by shifting resources to the miRNA biogenesis pathway at the onset of the dauer stage. Increasing the levels of a single pre-miRNA can have a global impact on the miRNA expression landscape; variations in rescue penetrance can then be attributed to the identity of the individual miRNA expressed and to the specific regulatory role it may hold (Wong et al., 2023).

To determine whether this transfer of information results in germline changes through the adjustment of gene expression via specific miRNA targets, we chose to use one of the miRNAs capable of suppressing the germline defects of AMPK mutants when expressed in the soma. *mir-34* is highly conserved and has been previously studied due to its localization to exosomes, its role in inter-tissue communication, and because of its identity as a tumour suppressor (Isik et al., 2016; Huang et al., 2020; Garcia-Martin et al., 2022). Potential *mir-34* mRNA targets were identified using TargetScanWorm (version 6.2) and were systematically subjected to RNAi in the *aak(0)* mutant background to determine if their reduction would bypass the need for the miRNA to ameliorate AMPK-dependent germline defects (Lewis et al., 2005; Jan et al., 2011). Reducing the expression of a number of these putative miRNA targets (thus mimicking the action of *mir-34*) was sufficient to partially restore germline integrity in the absence of AMPK signalling (Fig. 4F; Table S3). These miRNA targets may therefore be gene products that are misregulated in the absence of AMPK, thus contributing to the establishment of a gene expression profile inappropriate to the requirements of the dauer stage. In conclusion, we demonstrate that miRNAs produced downstream of AMPK signalling and of DCR-1 activity are necessary and sufficient to instruct germline stem cells to adapt their gene expression program during the dauer stage.

Because the somatic expression of a single miRNA, *mir-34*, is sufficient to partially restore germline integrity to animals that lack AMPK signalling, we sought to investigate if *mir-34*, or any miRNA capable of adjusting germline gene expression, was expressed in the germ cells in a dauer- and AMPK-dependent manner. We therefore established a means to visualize individual miRNAs and record their dynamics.

The small size of miRNAs makes them particularly challenging to image using commonly employed methods for RNA detection. To circumvent this matter, we established a protocol to adapt molecular beacon-based miRNA detection to visualize specific miRNAs in fixed *C. elegans* larvae. Molecular beacons are fluorophore- and quencher-conjugated probes which fold in the absence of a target and linearize upon binding to a complementary sequence, thus facilitating the highly sensitive, sequence-specific detection of mature miRNAs (Baker et al., 2012; Baker at al., 2013; James et al., 2017). Combining this method with widefield deconvolution microscopy, *mir-34*-specific probes were designed and used to detect *mir-34* both in the soma and in the germ cells of dauer larvae (Fig. 5A; Fig. S4). In contrast, no signal was detected in larvae that lack AMPK signalling, thus confirming the defects in miRNA production and the subsequent failure to signal to germline stem cells caused by the loss of AMPK at the onset of the dauer stage (Fig. 5A). We showed that expressing additional copies of DCR-1 could re-establish this equilibrium, presumably by making more DCR-1 available to associate with RBPL-1, thereby re-directing its activity to generate the appropriate miRNAs required for germline quiescence. Therefore, we reasoned that the additional DCR-1 activity could potentially restore the required levels of *mir-34* (and presumably of other miRNAs) in the germline stem cells. Consistent with this, driving the expression of *dcr-1* in the soma of AMPK mutants was sufficient to increase the abundance of *mir-34* in the soma and in the germ cells (Fig. 5B). These data complement our genetic analysis and provide strong evidence to demonstrate the importance of AMPK in re-allocating Dicer activity to promote the synthesis of pro-quiescence miRNAs involved in the establishment of germline stem cell quiescence at the onset of the dauer stage.

**Figure 5:**
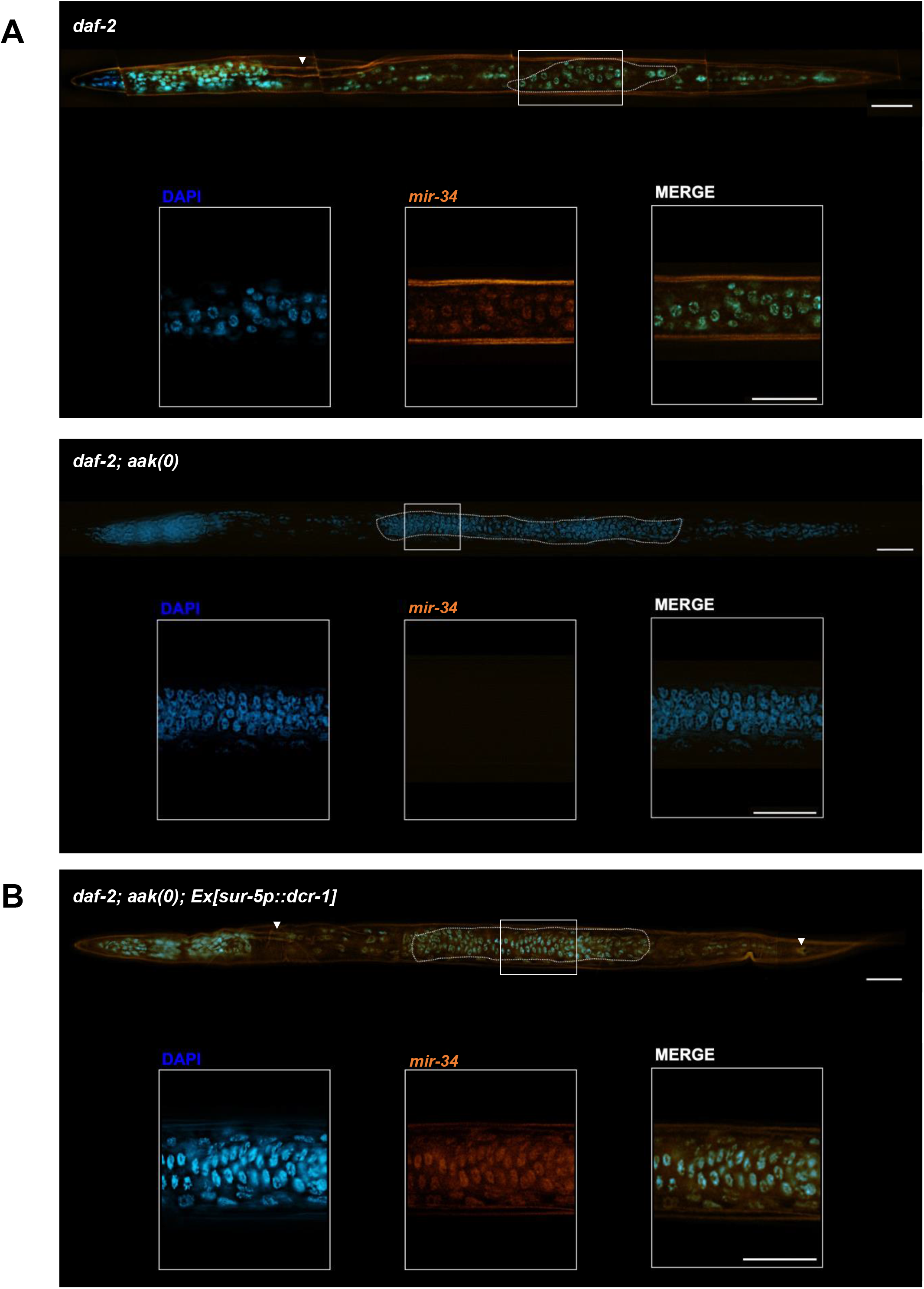
Production of a miRNA-derived signal in the germ cells of dauer larvae is under AMPK control. Animals were fixed and incubated with *mir-34*-complementary molecular beacons. The presence or absence of fluorescence was detected subsequent to processing and deconvolution of images representing (A) *daf-2* (top) and *aak(0)* (bottom) dauer larvae and (B) *aak(0)* dauer larvae expressing a *dcr-1* transgene (*aak(0); Ex[sur-5p::dcr-1])*. The hashed line delineates the boundary of the gonad. The white frame indicates the region of the gonad used to generate the high magnification insets. The arrowheads denote cells that exhibit neuronal morphology. Scale bar: 25 μM. All animals carry the *daf-2(e1370)* allele. See also Figure S4 for the imaging of a *mir-34* negative control using the same exposure parameters.

## Discussion

In recent years, several studies have provided data suggesting that the Weismann barrier – the unbridgeable gap separating soma and germ line – may not be as impermeable as was once believed (Conine & Rando, 2022; Devanapally et al., 2015; Dias & Ressler, 2014; Zeybel et al., 2012; Kadekar & Roy, 2019). Data from a number of laboratories using diverse models have demonstrated the role of inter-tissue communication and how it can drive gene expression changes within the germ line (Posner et al., 2019; Klosin et al., 2017; Cecere, 2021). These effects are often mediated by epigenetic modifications that manifest in the generation proper, or alternatively, in subsequent generations through transgenerational mechanisms that involve small RNAs. Here, we describe a mechanism which allows cells to differentially allocate resources for small RNA biogenesis to meet urgent cellular requirements for the production of one class of small RNA over another. These mechanisms occur in the soma and yet are critical to preserve germ cell integrity cell non-autonomously during periods of starvation-associated developmental arrest.

In response to environmental challenges the master regulator of metabolism AMPK targets factors that alter energy metabolism to ensure cellular energy homeostasis is maintained. However, in addition to phosphorylating cytoplasmic and mitochondrial metabolic enzymes, it also impinges on targets that directly and indirectly affect gene expression through both genetic and epigenetic mechanisms. The work we present here expands that toolbox by implicating AMPK in fine-tuning the equilibrium between the different cytoplasmic small RNA biosynthetic pathways to favour the production of critical miRNAs that are required to instruct germ cells to adopt a quiescent state during periods of energy stress.

Our small RNA analysis of AMPK mutant dauer larvae indicated that a major alteration in small RNA metabolism occurs in dauer larvae that lack AMPK signalling. Several small RNA classes are upregulated, while conversely, the miRNA class is globally downregulated. This reciprocal change in small RNA abundances hinted at the possible implication of a regulatory hingepin that acts upstream of all the affected pathways.

This global perturbation of small RNA homeostasis is likely the cause of the germline defects that arise in AMPK mutant dauer larvae, since compromising the activity of various small RNA regulators, and increasing the availability of DCR-1, both greatly improve germline function, while ultimately restoring post-dauer fertility in otherwise sterile animals. It remained unclear, however, if the sterility was a result of the increased abundance of small RNAs involved in endo-siRNA signalling, whether it was linked to the reduction of miRNAs, or a combination of both. To our surprise, expressing individual pre-miRNAs was sufficient to partially suppress the germline defects typical of the AMPK mutant dauer larvae. These data supported a major role for miRNAs in establishing germline quiescence during the dauer stage, in a manner that is independent of the other small RNA perturbations that occur. They further implicated that a putative AMPK target could affect the various branches of the small RNA pathways in a reciprocal manner.

Our understanding of how small RNAs contribute to cellular homeostasis has advanced rapidly since the discovery of the dsRNA trigger involved in exogenous RNAi. However, the manner by which the various pathways are regulated during development or during physiological challenges has not been systematically elucidated. Furthermore, although it was well appreciated that the major small RNA pathways rely on, and compete for, the endoribonuclease DCR-1, it was not clear how this enzyme could distinguish which branch to prioritize (Lee et al., 2006; Loinger et al., 2012; Zhuang & Hunter, 2012; Sawh & Duchaine, 2013). We show that AMPK impinges upon both DCR-1 and its pivotal binding partner RBPL-1, hence sequestering DCR-1 activity for the production of critical pro-quiescent miRNAs required to arrest germ cell proliferation and development.

Our data supports a model wherein DCR-1 differentially associates with various interactors to produce siRNAs and miRNAs in animals living in environments favourable for growth and reproduction. However, if animals are confronted with stressful growth conditions that require the conservation of energy and resources, AMPK promotes the association of RBPL-1, DCR-1, and pre-miRNA substrates. This interaction favours the production of a population of miRNAs in the soma, many of which are responsible for initiating the gene expression changes required for germ cells to adapt to the demands of a stressful growth environment (Fig. 6).

**Figure 6:**
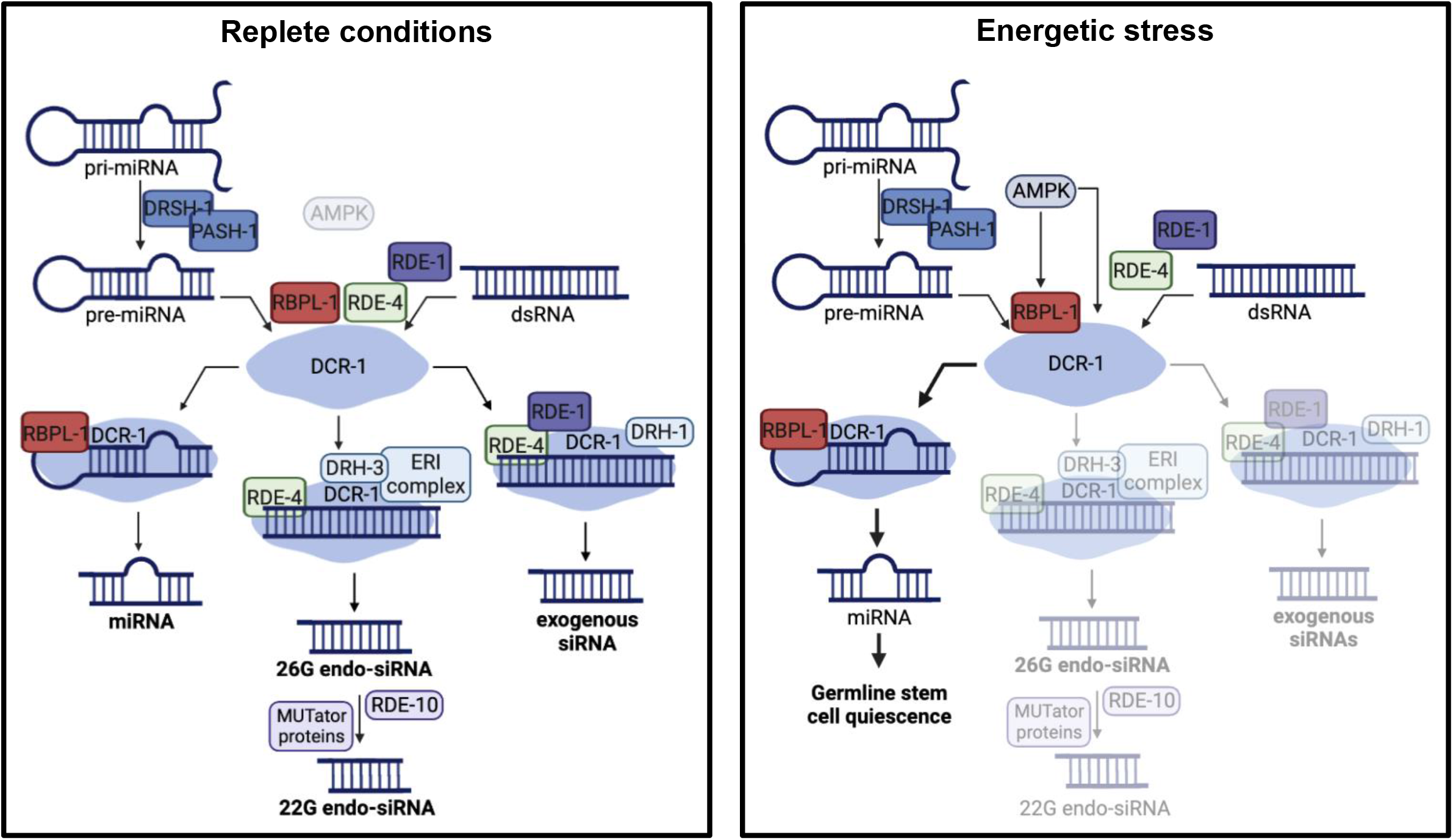
AMPK regulates Dicer-dependent small RNA production to favour the synthesis of pro-quiescent miRNAs for germline stem cell arrest. Throughout development, DCR-1 differentially associates with different siRNA and miRNA biogenesis factors to produce different types of small RNAs, as the requirement for them arises. Upon entry into the dauer stage, however, activated AMPK acts upon both RBPL-1 and DCR-1 to promote the formation of a complex containing DCR-1, RBPL-1 and the pre-miRNA substrate, thus favouring the generation of pro-quiescent miRNAs. This interaction is disrupted in the absence of AMPK signalling, causing the sequestration of DCR-1 to the siRNA biogenesis pathway and the impaired production of miRNAs required for the establishment of germline stem cell quiescence and the preservation of germline integrity.

We identified consensus phosphorylation sites on these important miRNA regulators (DCR-1 and RBPL-1) through bioinformatic prediction. The implication of these sites was validated by substituting the critical phospho-acceptor residues with phosphomimetic residues which partially suppressed the post-dauer sterility of AMPK mutant animals. Previous studies have demonstrated the existence of different protein isoforms of RBBP6, the human orthologue of RBPL-1 (Di Giammartino et al., 2014; Hull et al., 2015). Interestingly, further work also showed that the shortest isoform produced is often downregulated in human cancers (Mbita et al., 2012). Similarly, we demonstrate that hyperproliferation of germline stem cells caused by the loss of AMPK signalling correlates with a decreased interaction between Dicer and the short isoform of RBPL-1. In *C. elegans*, only the long isoform of RBPL-1 contains a putative AMPK phosphorylation site. We thus propose that AMPK acts on the long isoform of RBPL-1 at the onset of the dauer stage, ultimately driving the strengthened association of DCR-1 and of the short isoform of RBPL-1. The presence of a 14-3-3 binding site in the sequences of both isoforms of RBPL-1 could indeed further point to regulation by this protein, whose association with RBPL-1 would then depend upon RBPL-1 phosphorylation by AMPK (Morrison, 2009; Kundu et al., 2020).

Mammalian Dicer also harbours consensus phosphorylation sites, suggesting that AMPK may regulate this critical endonuclease in a similar manner to favour miRNA biogenesis under specific physiological conditions. Although the presence of phosphorylation sites does not necessarily coincide with phosphoregulation *in vivo*, several phosphoisoforms of Dicer have been identified in response to diverse experimental conditions and in a wide spectrum of tumour types (Burger et al., 2017; Cheng et al., 2017; Aryal et al., 2019). This would suggest that this endoribonuclease represents a conserved key regulatory target, typical of a hingepin component in the control of the flux of small RNAs required to respond to a multitude of physiological stimuli.

Curiously, we noted that a collection of cells that appear to be neurons (Fig. 5A-B) figure among the somatic cells that express *mir-34.* The role of these miRNAs in the neurons is intriguing, but is consistent with recent work which suggests that the miRNA pathway is critical for the AMPK-mediated adjustment that occurs in the germ cells at the onset of dauer (Wong et al. 2023). Although our work did not identify whether these pro-quiescence miRNAs are synthesized *de novo* or selected from an existing miRNA stockpile, future studies will clarify the pathways that are involved, and how the transmission of a small RNA-encoded pro-quiescence signal can instruct the germ cells to adapt their gene expression program accordingly. Our findings will help to define a previously undiscovered mechanism of RNA-based cellular communication and reveal the potential of this means of signalling for the development and delivery of novel RNA-based therapeutics.

## Acknowledgements

We thank all members of the Roy and Zetka labs for their comments and discussions. We thank Dr. Monique Zetka for allowing us to access her ZEISS Axio Observer microscope and for providing the anti-HA antibody. We also thank Ryan Dawson for technical assistance. Some strains were provided by the *C. elegans* Genetics Center (CGC), which is funded by NIH Office of Research Infrastructure Programs (P40 OD010440). This work was supported by the Canadian Institutes of Health Research (Project Grant, CIHR PJT-180267), the CIHR Banting and Best Doctoral Research Award, the University of Basel Excellent Junior Researcher Grant, the Swiss National Science Foundation (SNSF) SPARK Grant (CRSK-3_195955), the University of Toronto.

## Author Contributions

E.M.J. designed and carried out the experiments, analysed the collected data, wrote and edited the manuscript. C.W. aided in the design and execution of the experiments and edited the manuscript. F.B. and E.M. performed and analysed small RNA sequencing. T.F.D. and A.N.S. first characterised the regulation of Dicer by AMPK, provided materials and suggestions, and edited the manuscript. R.R. designed the experiments, wrote and edited the manuscript.

## Declaration of interests

The authors declare no competing interests.

## Methods

### *C. elegans* strains and maintenance

*C. elegans* strains were cultured at 15°C and grown on nematode growth media (NGM). Plates were seeded with *E. coli* bacteria (OP50) according to standard protocol (Brenner, 1974). Strains used in this study are listed in the Key Resources Table.

Transgenic lines and genetic mutants were generated according to standard molecular genetics approaches. Injection mixes were composed of: constructed plasmid (concentrations listed in Key Resources Table), pRF-4 (*rol-6*; 120 ng/µL), pSK(70 ng/µL), H_2_O.

### Plasmid Cloning and Preparation

Cloning and preparation of plasmid DNA were done according to Gibson Assembly Cloning Kit protocol and Biobasics: EZ-10 Spin Column Plasmid DNA Miniprep Kit protocol, respectively. The concentration of DNA was determined using Nanodrop 2000c Spectrophotometer 606 (Thermo Scientific).

The somatic overexpression of miRNAs was obtained by amplifying the individual pre-miRNA sequences and ∼1kb of downstream sequence from wild-type genomic DNA and cloning this fragment into a modified pPD95_77 vector containing the *sur-5p* promoter by Gibson assembly.

The tissue-specific overexpression of *dcr-1* was obtained by amplifying the sequence of *dcr-1* from wild-type genomic DNA and inserting it into a modified pPD95_77 vector by Gibson assembly. Tissue-specific promoters were amplified from wild-type genomic DNA and inserted in the *dcr-1*-containing vector by restriction digest and T4 DNA ligation.

RNAi clones not available in the Ahringer *C. elegans* RNAi library were made by amplifying a 600-1000 bp fragment of the target gene from wild-type cDNA and cloning these inserts into a modified pPD129.36 (L4440) vector by T4 DNA ligation. Plasmids were transformed into HT115 bacterial cells.

### Synchronization and Dauer Recovery Assay

*C. elegans* adults were synchronized using alkaline hypochlorite bleaching solution, as previously described (Porta-de-la-Riva et al., 2012). Worms were washed with distilled water and centrifuged for 40 seconds at 1.9 rpm. The supernatant was discarded, 1-2 mL of alkaline hypochlorite solution were added and the tube was vortexed for six minutes. The worm pellet was washed twice with distilled water and then resuspended before the embryos were plated on NGM or RNAi plates with a Pasteur pipette.

Animals were incubated at 25°C for 96 hours before being switched to the permissive temperature of 15°C to induce dauer recovery. Post-dauer fertility, rates of larval arrest, and somatic defects were assessed 7 days after the initiation of recovery.

### RNA interference

RNAi plates were made with NGM containing ampicillin and 1 mM IPTG. Mutants overexpressing *dcr-1* are RNAi hypersensitive, and as such were grown on RNAi plates containing 0.1 mM IPTG.

RNA interference was performed on standard feeding protocols as described elsewhere (Kamath et al., 2003). Briefly, RNAi clones were obtained from the Ahringer *C. elegans* RNAi feeding library (Fraser et al., 2000) and were cultured in LB medium supplemented with ampicillin at 37°C overnight. RNAi plates were seeded with individual clones and the bacteria were allowed to grow for at least 48 hours to induce the expression of dsRNA before embryos were synchronized on the plates. Animals were then made to traverse the dauer stage and to undergo dauer recovery on RNAi plates, as described above.

### Small RNA sequencing

RNA was isolated from wild-type and *aak(0)* dauer larvae via TRIzol extraction and resuspended in nuclease-free H_2_O.

Library preparation, RNA sequencing, and analysis of results were performed by Dr. Fabian Braukmann and Dr. Eric Miska. TruSeq Small RNA Library Preparation kits were used for library generation.

Clustering was performed calculating the Euclidian distance using the Matlab function “clustergram”. Read counts were log transformed.

### CRISPR-Cas9 Genome Editing

CRISPR-Cas9 genome editing was used to generate DCR-1 and RBPL-1 phosphomimetic strains. DCR-1(T961D) was generated using the sgRNA site CGTCGTTCAAGAACTGTGAG. The threonine residue was mutated to an aspartic acid (ACT→GAT) using the repair template listed in the Key Resources Table. Homozygous DCR-1(T961D) was found to not be viable; as such, animals were maintained as heterozygotes (DCR-1(T961D)/+). RBPL-1(S367E) was generated using the sgRNA site GAGCACTAAGTGATGTCCC. The serine residue was mutated to a glutamic acid (TCA→GAA) using the repair template listed in the Key Resources Table.

An HA tag (TACCCATACGATGTTCCAGATTACGCT) was added to the C-terminus of RBPL-1 and of RDE-4 using the sgRNAs TTGGAACTTTTCGAGACGA and TTTCAACACCTATGATTTC, respectively. A FLAG tag (GACTACAAAGACGATGAC GACAAG) was added to the C-terminus of DCR-1 using the sgRNA CTTCACTTTCTGTGATATGCAAGACATCTCGCAATAG. The repair templates used are listed in the Key Resources Table.

sgRNAs were individually cloned into the pDD162 vector (Peft-3::Cas9 + Empty sgRNA).

Injection mixes were composed of: repair template (40 ng/µL), sgRNA (60 ng/µL), pRF-4 (*rol-6*; 120 ng/µL), H_2_O.

### Western Blot analysis

Western Blot analysis was conducted as described elsewhere (Kadekar & Roy, 2019). 500 dauer larvae or 200 post-dauer adults were picked and resuspended into 20 or 40 µL of PBST, respectively. 5x SDS loading buffer (5% β-mercaptoethanol, 0.02% bromophenol blue, 30% glycerol, 10% sodium dodecyl sulfate, 250 mM pH 6.8 Tris-Cl) was added and the samples underwent 3-5 freeze-thaw cycles of 3 minutes each (alternating between liquid nitrogen and 100°C heat block), before being loaded on a 6% polyacrylamide gel in SDS running buffer. Proteins were transferred onto a nitrocellulose membrane. The membrane was incubated with rabbit anti-FLAG, mouse anti-HA, or mouse anti-α-tubulin. After SDS-PAGE and Western blotting, proteins were visualized using horseradish-peroxidase-conjugated anti-rabbit or anti-mouse secondary antibodies. The signal was developed using Clarity Western ECL Substrate and visualized using MicroChemi (DNR Bio Imaging Systems) and GelCapture software (Version 7.0.18).

### Co-immunoprecipitation

Co-immunoprecipitation against the FLAG tag and the HA tag was performed using the Abcam Immunoprecipitation protocol adapted for populations of dauer larvae. Protein A beads were used according to the manufacturer’s instructions. Input and IP were loaded onto an SDS-PAGE gel and Western Blotting was performed as described above. The primary antibodies used were anti-FLAG and anti-HA.

### Identification of putative AMPK phosphorylation motifs

The protein sequences of DCR-1 and of known DCR-1 interactors were obtained through Wormbase and input into Scansite, version 4.0. The sequences were then scanned for short protein motifs recognized by AMPK with high, medium, or low stringency.

### RT-qPCR

RNA was isolated from wild-type, *aak(0)*, and *aak(0); Ex[sur-5p::mir-51]* dauer larvae via TRIzol extraction and resuspended in nuclease-free H_2_O. The reverse transcription and qPCR reactions were performed using the Taqman MicroRNA Reverse Transcription Kit, a *mir-235* Taqman MicroRNA Assay, and the Taqman Universal Master Mix II, with UNG. qPCR reactions were performed using the CFX96 Touch Real-Time PCR Detection System and the CFX Manager 3.0 Software.

### miRNA detection in *C. elegans* dauer larvae

A *mir-34*-specific molecular beacon was designed following guidelines described by Baker et al., 2013 (sequence listed in Key Resources Table). This beacon was designed using a DNA backbone, which exhibits higher affinity for mature miRNAs compared to pre-miRNAs and pri-miRNAs (Baker et al., 2012; Baker et al., 2013).

Fixation and hybridization of the probes followed adapted *C. elegans* smFISH methodology (Ji N. & van Oudenaarden A., 2012; Paramasivam et al., 2022). Animals were made to transit through the dauer stage as described above, and were resuspended in M9 buffer before being fixed for 30 min – 2 hours in 1mL of fixation buffer (7.4% PFA in 1x PBS). The worms were washed twice with 1 mL of 1x PBS, then were resuspended and kept rotating overnight in Carnoy’s solution (60% ethanol, 30% chloroform, 10% acetic acid). Hybridization solution was made by adding the probe at a concentration of 10 nM to hybridization buffer (10% dextran sulfate, 0.1% *E. coli* tRNA, 100 µL 200 mM vanadyl ribonucleoside complex, 40 µL 50 mg/mL RNase-free BSA, 10% formamide, nuclease-free H_2_O to 10 mL). The fixed animals were resuspended in 1 mL wash buffer (10% formamide, 10% 20x SSC, 80% nuclease-free H_2_O); the sample was centrifuged and the wash buffer aspirated. The hybridization solution was added and samples were incubated overnight at 30°C in the dark. Samples were subjected to two 30-min incubations in wash buffer at 30°C. DAPI stain was added during the second incubation. Samples were finally washed and resuspended in 2x SSC, and were imaged immediately after hybridization using the ZEISS Axio Observer LED fluorescence microscopy. Deconvolution (settings: constrained iterative and Clip normalization, factor 1.00) and stitching were applied during image processing through the software ZEISS Zen 3.7.

## Notes

### Competing Interest Statement

The authors have declared no competing interest.

