## Supplementary Figures and Tables for "AMPK regulates small RNA pathway prevalence to mediate soma-to-germ line communication and establish germline stem cell quiescence"

| **Protein analysed** | **Number of predicted AMPK phosphorylation motif sites** | **Motif stringency** |
|---|---|---|
| **DCR-1** | **13** | **Low (2), medium (11), high (1)** |
| ERI-1 | 1 | Low (1) |
| ERI-3 | 0 | N/A |
| ERI-5 | 0 | N/A |
| ERI-6 | 2 | Low (1), medium (1) |
| ERI-9 | 5 | Low (5) |
| RDE-4 | 4 | Low (3), medium (1) |
| HENN-1 | 1 | Low (1) |
| DRH-3 | 6 | Low (4), medium (2) |
| EGO-1 | 11 | Low (8), medium (2), high (1) |
| EKL-1 | 2 | Low (2) |
| MUT-14 | 0 | N/A |
| MUT-15 | 5 | Low (3), medium (1), high (1) |
| RDE-10 | 4 | Low (3), medium (1) |
| RDE-11 | 1 | Low (1) |
| DRSH-1 | 7 | Low (6), medium (1) |
| PASH-1 | 5 | Low (3), medium (2) |
| ALG-1 | 8 | Low (6), medium (2) |
| ALG-2 | 2 | Low (2) |
| AIN-1 | 4 | Low (3), medium (1) |
| AIN-2 | 7 | Low (5), medium (2) |
| **RBPL-1** | **19** | **Low (13), medium (5), high (1)** |

**Supplemental Table 1**
The protein sequences of DCR-1 and of DCR-1 interactors (including siRNA and miRNA biogenesis factors, co-factors, and RISC components) were analysed for the presence of low, medium, and high stringency putative AMPK phosphorylation motifs using the software ScanSite 4.0. RBPL-1 and DCR-1 (in bold) were thus identified as potential targets of AMPK phosphorylation and the putative phosphorylation sites on these proteins were subsequently subjected to further analyses. See also Fig. 4B.

| **miRNA family** | **Members** | **Members downregulated in AMPK mutants** |
|---|---|---|
| *let-7* | *let-7, mir-48, mir-84, mir-241, mir-265, mir-795, mir-1821, mir-793, mir-794* | *let-7, mir-48, mir-84, mir-241, mir-793, mir-794* |
| *mir-2* | *mir-2, mir-43, mir-250, mir-797* | *mir-2, mir-250* |
| *mir-229* | *mir-229, mir-63, mir-64, mir-65, mir-66* | *mir-229, mir-63, mir-64, mir-65, mir-66* |
| *mir-35* | *mir-35, mir-36, mir-37, mir-38, mir-39, mir-40, mir-41, mir-42* | *mir-38* |
| *mir-44* | *mir-44, mir-45, mir-61, mir-247* | *mir-44, mir-61* |
| *mir-51* | *mir-51, mir-52, mir-53, mir-54, mir-55, mir-56* | *mir-51, mir-53, mir-54, mir-55, mir-56* |
| *mir-58* | *mir-58, mir-80, mir-81, mir-82, mir-1834* | *mir-58, mir-81, mir-82* |
| *mir-72* | *mir-72, mir-73, mir-74* | *mir-72, mir-73* |
| *mir-49* | *mir-49, mir-83* | *mir-83* |
| *mir-34* | *mir-34, mir-1824* | *mir-34* |

**Supplemental Table 2**

A fraction of miRNAs identified through small RNA sequencing as being downregulated in AMPK mutant dauers were categorized according to the family to which they belong, as previously defined through literature (Alvarez-Saavedra & Horvitz, 2010). This analysis permitted the identification of highly conserved miRNAs that were selected for transgenic expression in *aak(0)* mutants. See also Fig. 4D.

| **mRNA targets with no significant effect on *aak(0)* post-dauer fertility** | | |
|---|---|---|
| C44C10.9 | K11H12.7 | *larp-5* |
| *hlh-11* | F45E4.5 | C37C3.6 |
| *lmn-1* | F07C3.2 | *twf-2* |
| *myo-5* | *sams-1* | *vab-10* |
| *unc-52* | *utx-1* | *let-805* |
| *htz-1* | C09F5.1 | *lir-1* |
| C02B10.3 | C16D9.1 | Y97E10C.1 |
| *lec-3* | *col-181* | *elf-1* |
| *egl-44* | *slfl-5* | *ost-1* |
| *zyx-1* | *emb-9* | T23B12.11 |
| C26G2.2 | ZK1248.13 | K08D12.6 |
| *col-101* | *ulp-4* | *cpg-9* |
| *sem-5* | *dep-1* | C50F2.8 |
| *lgg-2* | *mtm-3* | F42G8.10 |
| F11E6.3 | *unc-70* | W03G11.4 |
| T21B6.3 | F32B5.6 | *deb-1* |
| *ZK632.9* | *gon-1* | *unc-64* |
| *gsnl-1* | *cle-1* | D2005.6 |
| *lim-7* | D2023.1 | |
| *pqn-52* | *cerk-1* | |

**Supplemental Table 3**

The software TargetScanWorm (Version 6.2) was used to identify putative mRNA targets of *mir-34* by searching for the presence of the *mir-34* seed sequence in the 3'UTR. An RNAi survey was performed to evaluate the effects of individual genes in the *aak(0)* mutant background; animals were induced to form, and allowed to transit through dauer, after which post-dauer fertility was assessed. See also Fig. 4F.

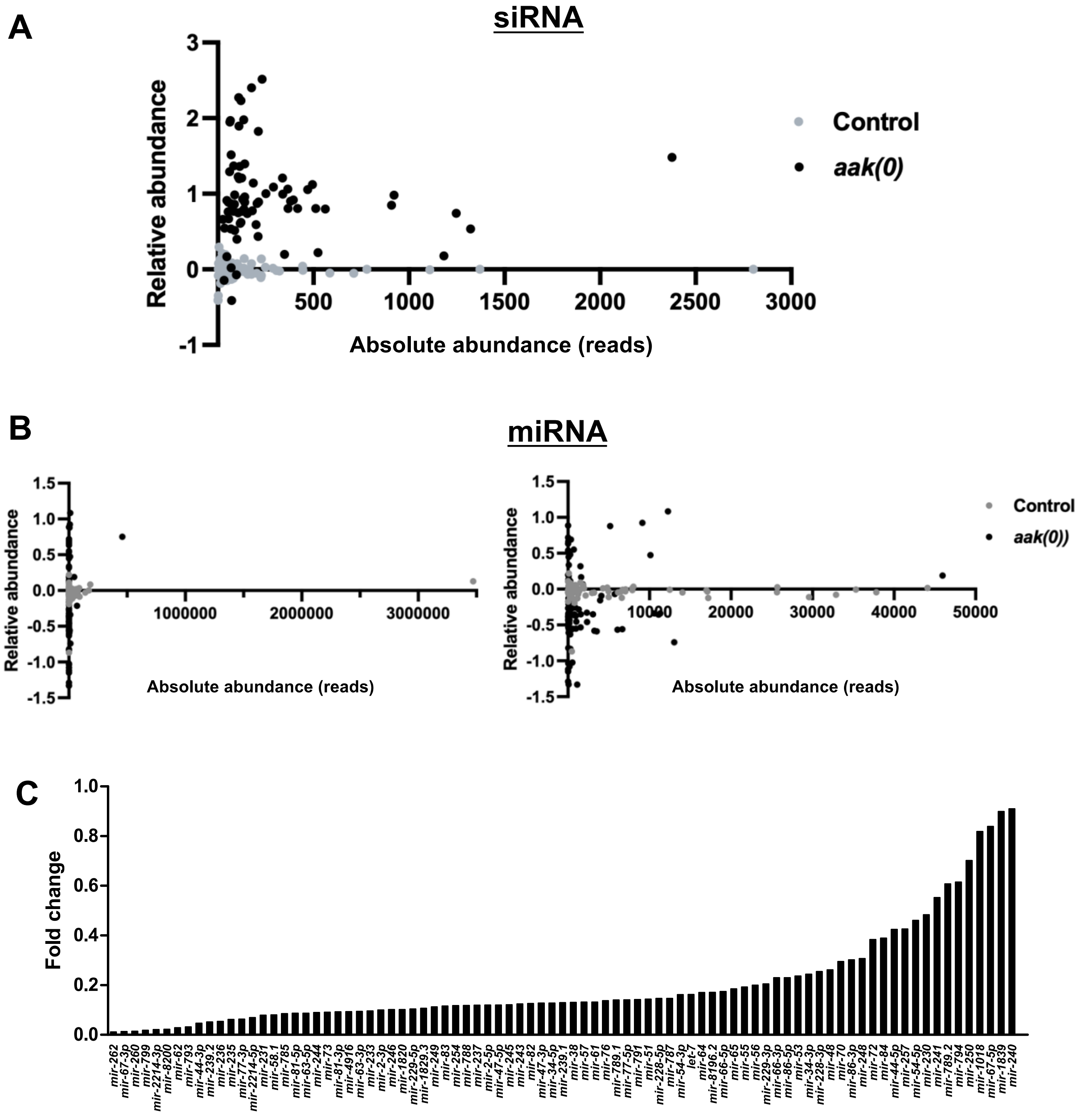


**Supplemental Figure 1**

Following small RNA sequencing analysis, changes in siRNA and miRNA expression levels in control and *aak(0)* larvae that recovered from the dauer stage were plotted. (A-B). Relative vs absolute abundance plots of reads per gene for siRNAs and miRNAs in wild-type and *aak(0)* background. (C) Graphical representation of miRNA downregulation in *aak(0)* dauer mutants. miRNAs are ranked based on the degree of fold change in expression.

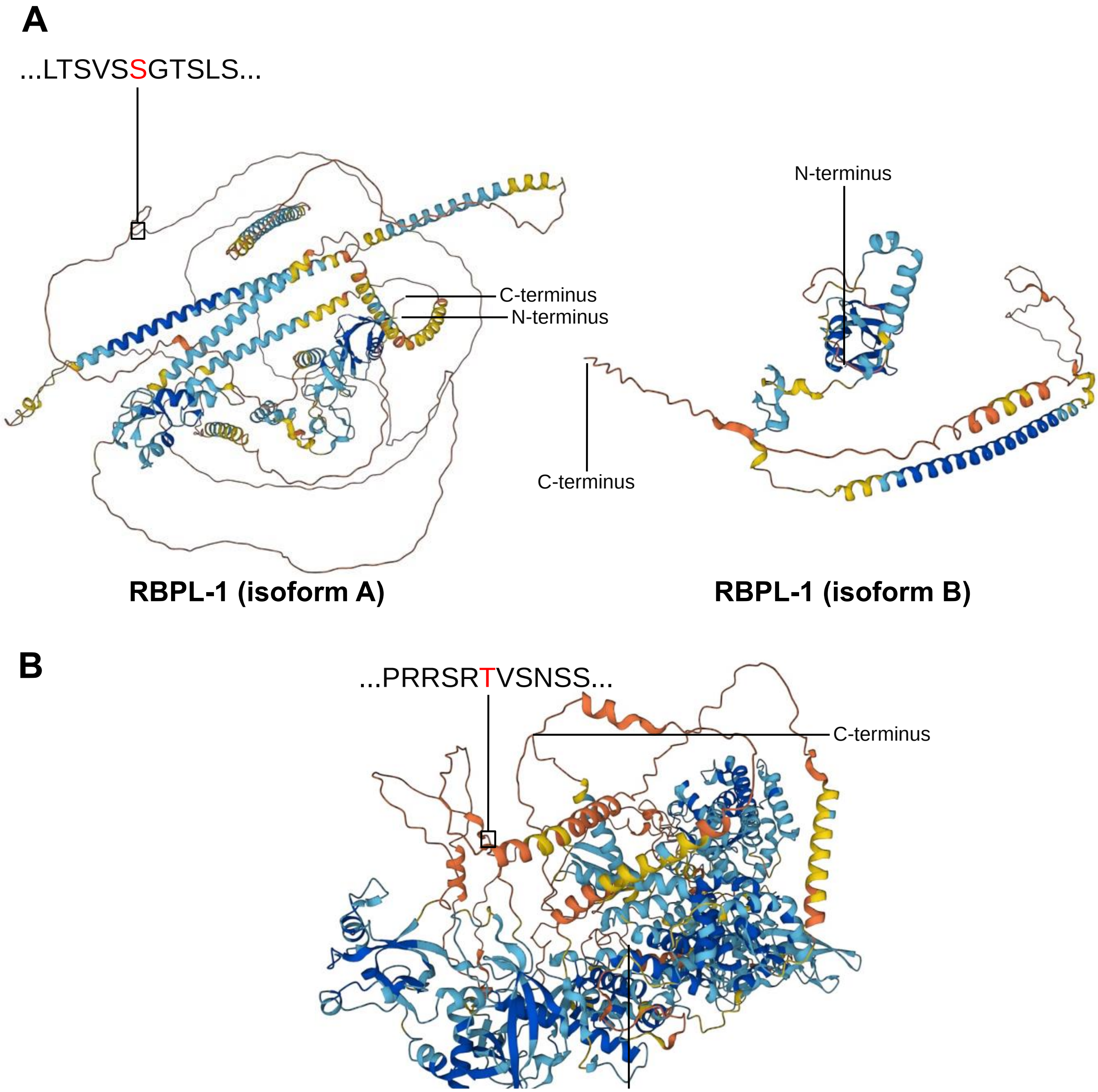


**Supplemental Figure 2**

The sequences of RBPL-1 and of DCR-1 were scanned for the presence of putative high stringency AMPK phosphorylation motifs and protein plots were generated using the software ScanSite 4.0. This analysis permitted the identification of an LxSxxS motif at position 362-367 on RBPL-1 and of a RXXT motif at position 958-961 on DCR-1 and the subsequent generation of phosphomimetic mutations at these sites. Predicted protein structures of (A) both isoforms of RBPL-1 and (B) DCR-1 (full-length protein) were produced by AlphaFold Protein Structure Database. See also Fig. 4B-C and Fig. S3.

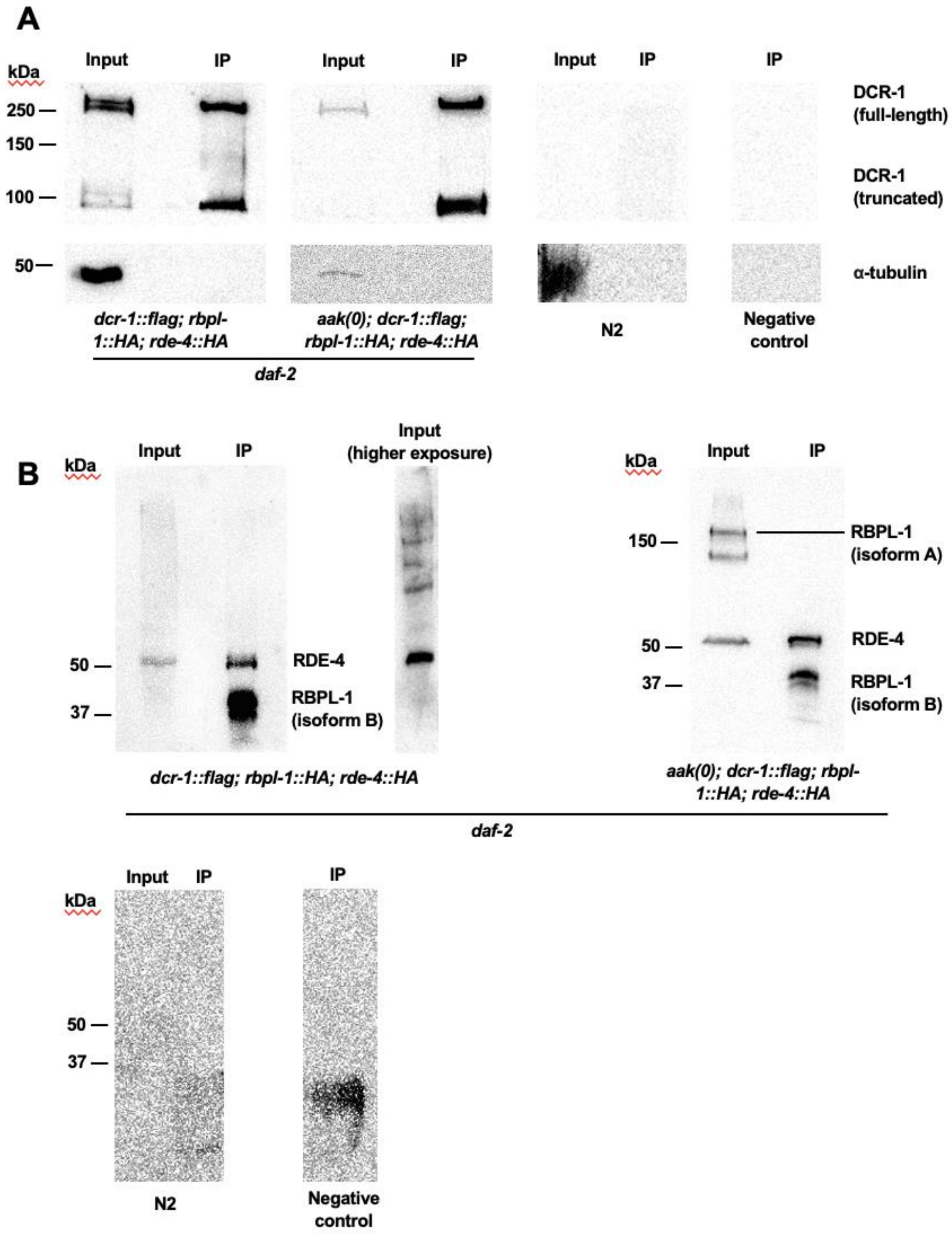


**Supplemental Figure 3**

Immunoprecipitation using an antibody against the FLAG tag was performed to immunoprecipitate DCR-1::FLAG both in control and *aak(0)* dauer larvae. Two negative controls are shown: IP performed using N2 animals and IP performed using no protein lysate (denoted as “Negative control”). (A) IP membranes were probed with an anti-FLAG primary antibody to detect the full-length and truncated forms of DCR-1. (B) IP membranes were probed with an anti-HA primary antibody, permitting the detection of RDE-4::HA and of RBPL-1::HA

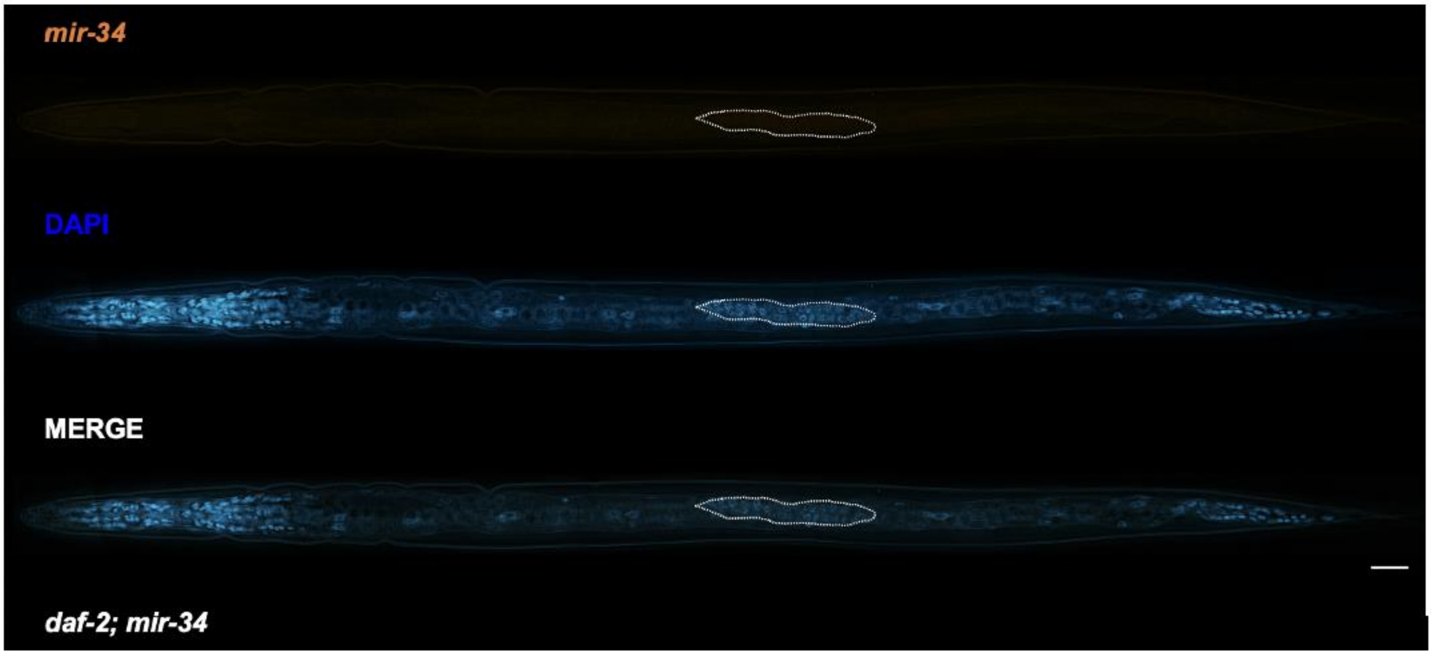


**Supplemental Figure 4**
Dauer larvae of *mir-34* mutants were fixed and incubated with *mir-34*-complementary molecular beacons. No signal is detected when appropriate exposure signals are applied. The high sensitivity and specificity of molecular beacons combined with deconvolution microscopy thus provides optimal detection of signal and low background fluorescence. The hashed line delineates the gonad boundary. Scale bar: 25 μM. Animals carry the *daf-2(e1370)* allele.
