## Supplementary Information for "AMPK regulates small RNA pathway prevalence to mediate soma-to-germ line communication and establish germline stem cell quiescence"

**Experimental models: *C. elegans* strains**

| Strain | Genotype | Source |
|---|---|---|
| N2 | *C. elegans* wild isolate | CGC |
| CB1370 | *daf-2(e1370)III* | CGC |
| MR1000 | *daf-2(e1370) aak-1(tm1944)III; aak-2(ok523)X* | Roy Laboratory |
| MR1737 | *rrf-3(pk1426)II; daf-2(e1370)III* | Roy Laboratory |
| MR1689 | *daf-2(e1370)III; rde-1(ne219)V* | Roy Laboratory |
| MR1963 | *daf-2(e1370) aak-1(tm1944)III; tbc-7(rr166) aak-2(ok523)X* | Roy Laboratory |
| MR2369 | *daf-2(e1370) aak-1(tm1944)III; aak-2(ok524)X; rrEx499[sur-5p::dcr-1::gfp;rol-6].* Line 1. 10 ng/µL. | This study |
| MR2370 | *daf-2(e1370) aak-1(tm1944)III; aak-2(ok524)X; rrEx500[sur-5p::dcr-1::gfp;rol-6]* Line 2. 10 ng/µL. | This study |
| MR2371 | *daf-2(e1370) aak-1(tm1944)III; aak-2(ok524)X; rrEx501[sur-5p::dcr-1::gfp;rol-6]* Line 3. 10 ng/µL. | This study |
| MR2373 | *daf-2(e1370) aak-1(tm1944)III; aak-2(ok524)X; rrEx503[sur-5p::dcr-1::gfp;rol-6].* 1 ng/µL. | This study |
| MR2483 | *daf-2(e1370) aak-1(tm1944)III; aak-2(ok524)X; rrEx561[rgef-1p::dcr-1; rol-6]*. Line 1. 10 ng/µL. | This study |
| MR2886 | *daf-2(e1370) aak-1(tm1944)III; aak-2(ok524)X; rrEx887[rgef-1p::dcr-1; rol-6]*. Line 2. 10 ng/µL. | This study |
| MR2887 | *daf-2(e1370) aak-1(tm1944)III; aak-2(ok524)X; rrEx888[rgef-1p::dcr-1; rol-6]*. Line 3. 10 ng/µL. | This study |
| MR2426 | *daf-2(e1370) aak-1(tm1944)III; aak-2(ok524)X; rrEx508[pgp-12p::dcr-1; rol-6]*. Line 1. 10 ng/µL. | This study |
| MR2834 | *daf-2(e1370) aak-1(tm1944)III; aak-2(ok524)X; rrEx862[pgp-12p::dcr-1; rol-6]*. Line 2. 10 ng/µL. | This study |
| MR2835 | *daf-2(e1370) aak-1(tm1944)III; aak-2(ok524)X; rrEx863[pgp-12p::dcr-1; rol-6]*. Line 3. 10 ng/µL. | This study |
| MR2488 | *daf-2(e1370) aak-1(tm1944)III; aak-2(ok524)X; rrEx566[wrt-2p::dcr-1; rol-6]*. Line 1. 10 ng/µL. | This study |
| MR2828 | *daf-2(e1370) aak-1(tm1944)III; aak-2(ok524)X; rrEx856[wrt-2p::dcr-1; rol-6]*. Line 2. 10 ng/µL. | This study |

| | | |
|---|---|---|
| MR2829 | *daf-2(e1370) aak-1(tm1944)III; aak-2(ok524)X; rrEx857[wrt-2p::dcr-1; rol-6]*. Line 3. 10 ng/µL. | This study |
| MR2482 | *daf-2(e1370) aak-1(tm1944)III; aak-2(ok524)X; rrEx560[myo-3p::dcr-1; rol-6]*. Line 1. 10 ng/µL. | This study |
| MR2830 | *daf-2(e1370) aak-1(tm1944)III; aak-2(ok524)X; rrEx858[myo-3p::dcr-1; rol-6]*. Line 2. 10 ng/µL. | This study |
| MR2831 | *daf-2(e1370) aak-1(tm1944)III; aak-2(ok524)X; rrEx859[myo-3p::dcr-1; rol-6]*. Line 3. 10 ng/µL. | This study |
| MR2481 | *daf-2(e1370) aak-1(tm1944)III; aak-2(ok524)X; rrEx559[nhx-2p::dcr-1; rol-6]*. Line 1. 10 ng/µL. | This study |
| MR2832 | *daf-2(e1370) aak-1(tm1944)III; aak-2(ok524)X; rrEx860[nhx-2p::dcr-1; rol-6]*. Line 2. 10 ng/µL. | This study |
| MR2833 | *daf-2(e1370) aak-1(tm1944)III; aak-2(ok524)X; rrEx861[nhx-2p::dcr-1; rol-6]*. Line 3. 10 ng/µL. | This study |
| MR2409 | *daf-2(e1370) aak-1(tm1944)III; eri-1(ok2683)IV; aak-2(ok524)X* | This study |
| MR2374 | *eri-3(tm1361)II; daf-2(e1370) aak-1(tm1944)III; aak-2(ok524)X* | This study |
| MR2423 | *eri-6(mg379)I; daf-2(e1370) aak-1(tm1944)III; aak-2(ok524)X* | This study |
| MR2424 | *daf-2(e1370) aak-1(tm1944) eri-9(gg106)III; aak-2(ok524)X* | This study |
| MR2616 | *rbpl-1(S367E) I; daf-2(e1370) aak-1(tm1944)III; aak-2(ok524)X* | This study |
| MR2748 | *daf-2(e1370) aak-1(tm1944) dcr-1(T961D)III; aak-2(ok524)X* | This study |
| MR2849 | *rbpl-1(rr184[rbpl-1::HA])I; daf-2(e1370) dcr-1(rr183[dcr-1::flag]) rde-4(rr185[rde-4::HA])III* | This study |
| MR2857 | *rbpl-1(rr184[rbpl-1::HA])I; daf-2(e1370) dcr-1(rr183[dcr-1::flag]) aak-1(rr189) rde-4(rr185[rde-4::HA])III; aak-2(ok524)X* | This study |
| MR2494 | *daf-2(e1370) aak-1(tm1944)III; aak-2(ok524)X; rrEx568[sur-5p::mir-1; rol-6]*. Line 1. 1 ng/µL. | Roy Laboratory |
| MR2495 | *daf-2(e1370) aak-1(tm1944)III; aak-2(ok524)X; rrEx569[sur-5p::mir-1; rol-6]*. Line 2. 1 ng/µL. | This study |
| MR2496 | *daf-2(e1370) aak-1(tm1944)III; aak-2(ok524)X; rrEx570[sur-5p::mir-1; rol-6]*. Line 3. 1 ng/µL. | This study |
| MR2564 | *daf-2(e1370) aak-1(tm1944)III; aak-2(ok524)X; rrEx627[sur-5p::mir-34; rol-6].* Line 1. 5 ng/µL. | This study |
| MR2565 | *daf-2(e1370) aak-1(tm1944)III; aak-2(ok524)X; rrEx628[sur-5p::mir-34; rol-6].* Line 2. 5 ng/µL. | This study |

| | | |
|---|---|---|
| MR2566 | *daf-2(e1370) aak-1(tm1944)III; aak-2(ok524)X; rrEx629[sur-5p::mir-34; rol-6].* Line 3. 5 ng/µL. | This study |
| MR2538 | *daf-2(e1370) aak-1(tm1944)III; aak-2(ok524)X; rrEx621[sur-5p::mir-44; rol-6].* Line 1. 10 ng/µL. | This study |
| MR2539 | *daf-2(e1370) aak-1(tm1944)III; aak-2(ok524)X; rrEx622[sur-5p::mir-44; rol-6].* Line 2. 10 ng/µL. | This study |
| MR2540 | *daf-2(e1370) aak-1(tm1944)III; aak-2(ok524)X; rrEx622[sur-5p::mir-44; rol-6].* Line 3. 10 ng/µL. | This study |
| MR2625 | *daf-2(e1370) aak-1(tm1944)III; aak-2(ok524)X; rrEx663[sur-5p::mir-51; rol-6].* Line 1. 10 ng/µL. | This study |
| MR2810 | *daf-2(e1370) aak-1(tm1944)III; aak-2(ok524)X; rrEx844[sur-5p::mir-51; rol-6].* Line 2. 10 ng/µL. | This study |
| MR2825 | *daf-2(e1370) aak-1(tm1944)III; aak-2(ok524)X; rrEx853[sur-5p::mir-51; rol-6].* Line 3. 10 ng/µL. | This study |
| MR2485 | *daf-2(e1370) aak-1(tm1944)III; aak-2(ok524)X; rrEx563[sur-5p::mir-58; rol-6]*. Line 1. 10 ng/µL. | This study |
| MR2486 | *daf-2(e1370) aak-1(tm1944)III; aak-2(ok524)X; rrEx564[sur-5p::mir-58; rol-6]*. Line 2. 10 ng/µL. | This study |
| MR2487 | *daf-2(e1370) aak-1(tm1944)III; aak-2(ok524)X; rrEx565[sur-5p::mir-58; rol-6]*. Line 3. 10 ng/µL. | This study |
| MR2569 | *daf-2(e1370) aak-1(tm1944)III; aak-2(ok524)X; rrEx631[sur-5p::mir-38; rol-6]*. Line 1. 10 ng/µL. | This study |
| MR2889 | *daf-2(e1370) aak-1(tm1944)III; aak-2(ok524)X; rrEx889[sur-5p::mir-38; rol-6]*. Line 2. 10 ng/µL. | This study |
| MR2890 | *daf-2(e1370) aak-1(tm1944)III; aak-2(ok524)X; rrEx890[sur-5p::mir-38; rol-6]*. Line 3. 10 ng/µL. | This study |
| MR2568 | *daf-2(e1370) aak-1(tm1944)III; aak-2(ok524)X; rrEx631[sur-5p::mir-84; rol-6]*. Line 1. 10 ng/µL. | This study |
| MR2920 | *daf-2(e1370) aak-1(tm1944)III; aak-2(ok524)X; rrEx911[sur-5p::mir-84; rol-6]*. Line 2. 10 ng/µL. | This study |
| MR2929 | *daf-2(e1370) aak-1(tm1944)III; aak-2(ok524)X; rrEx915[sur-5p::mir-84; rol-6]*. Line 3. 10 ng/µL. | This study |
| MR2885 | *daf-2(e1370)III; mir-34(gk437)X* | This study |

**Bacterial strains / source / identifier**

| | | |
|---|---|---|
| *Escherichia coli*: OP50 | CGC | N/A |
| *Escherichia coli*: HT115 | CGC | N/A |
| *Escherichia coli*: HB101 | CGC | N/A |

**Reagent or resource / source / identifier**

| | | |
|---|---|---|
| Protein A-Agarose | Sigma-Aldrich | P1406 |
| Halt™ Protease Inhibitor Cocktail (100X) | ThermoFisher Scientific | PI87786 |
| Ampicillin Sodium Salt | ThermoFisher Scientific | BP1760-25 |
| IPTG, ultra pure, Dioxane free | BioShop | IPT001 |
| TRIzol Reagent | ThermoFisher Scientific | 15596026 |
| DAPI | Roche | 10236276001 |

**Antibodies / source / identifier**

| | | |
|---|---|---|
| Anti-FLAG rabbit antibody | Sigma-Aldrich | F7425 |
| Anti-HA.11 epitope tag antibody | BioLegend | 901501 |
| HRP-conjugated goat anti-rabbit and goat anti-mouse secondary antibodies | Bio-Rad | 1706515<br>1706516 |

**Chemicals, peptides, and recombinant proteins / source / identifier**

| | | |
|---|---|---|
| Clarity Western ECL Substrate | Bio-Rad | 1705061 |
| Nitrocellulose membranes, 0.45 µm | Bio-Rad | 1620115 |

**Critical commercial assays / source / identifier**

| | | |
|---|---|---|
| TruSeq Small RNA Library Preparation Kit | Illumina | N/A |
| Taqman MicroRNA Reverse Transcription Kit | ThermoFisher Scientific | 4366596 |
| Taqman MicroRNA Assay | ThermoFisher Scientific | 4427975 |

| | | |
|---|---|---|
| Taqman Universal Master Mix II, with UNG | ThermoFisher Scientific | 4440042 |

**Oligonucleotides / source / identifier**

| | | |
|---|---|---|
| *dcr-1* primers used for Gibson assembly<br><br>Fragment 1:<br>5' ATTCAAAAGAGTCGAATACTTCCG 3'<br>5' TCCTCAAGCATCTGCTTCTG 3'<br><br>Fragment 2:<br>5' ATTCAAAAGAGTCGAATACTTCCG 3'<br>5' TTTGGGTCCTTTGGCCAATTAAACAGTTGTTAATGATGGGCTTT 3' | This study | IDT |
| *rgef-1p* primers (with SphI and XmaI restriction sites)<br>5' CATGGCATGCATCCTTTTCATTTTGAACTCACC 3'<br>5' CATGCCCGGGCGTCGTCGTCGTCGATGC 3' | This study | IDT |
| *pgp-12p* (with SphI and XmaI restriction sites)<br>5' CATGGCATGCTGGTGTGCGTGAGAAATCAT 3'<br>5' CATGCCCGGGGTTTAACCTATTTCAGAAGAATATCTGTTG 3' | This study | IDT |
| *myo-3p* (with SphI and XmaI restriction sites)<br>5' CATGGCATGCACAGTTCCAATTGCTACCGC 3'<br>5' CATGCCCGGGTTCTAGATGGATCTAGTGGTCGTG 3' | This study | IDT |
| *nhx-2p* (with SphI and XmaI restriction sites)<br>5' CATGGCATGCGACCACCACGTGTCTACTGA 3'<br>5' CATGCCCGGGCCAAGAATCAAAACAACCGGAAATAG 3' | This study | IDT |
| *wrt-2p* (with SphI and XmaI restriction sites)<br>5' CATGGCATGCGGTGTTCCCCATTCGAAAATAGT 3'<br>5' CATGCCCGGGCCGAGAAACAATTGGCAGGT 3' | This study | IDT |
| *sur-5p* (with SphI and XmaI restriction sites)<br>5' CATGGCATGCTTTTTGCGAAAGCCTACGAT 3' | This study | IDT |

| | | |
|---|---|---|
| 3’<br>CATGCCCGGGTCTGAAAACAAAATGTAAAGTTCAAAG 5’ | | |
| Primers used to generate *sur-5p::rbpl-1::HA*<br>5’<br>CTTTACATTTTGTTTTCAGAATGTCGTCAATTCACTACAAGT 3’<br>5’<br>TCTGGAACATCGTATGGGTACAAAAGATCAAATTTAATCTTGGTACTC 3’ | This study | IDT |
| Primers used to amplify microRNAs<br>*mir-1*<br>5’<br>CTTTACATTTTGTTTTCAGAAAAGTGACCGTACCGAGCT 3’<br>5’<br>CAGTTGGAATTCTACGAATGCATTGCCACGTCACAGAACT 3’<br>*mir-34*<br>5’<br>CTTTACATTTTGTTTTCAGACGGACAATGCTCGAGAGG 3’<br>5’<br>CAGTTGGAATTCTACGAATGAGCGTTTTAAAGAAGCGTCGA 3’<br>*mir-38*<br>5’<br>CTTTACATTTTGTTTTCAGAGTGAGCCAGGTCCTGTTC 3’<br>5’<br>CAGTTGGAATTCTACGAATGGCGGGTCAATTTCAGCTGAA 3’<br>*mir-44*<br>5’<br>CTTTACATTTTGTTTTCAGAGAGAAAATGGCCAATCTGGATG 3’<br>5’<br>CAGTTGGAATTCTACGAATGGCGGTGTAACAGGGTCAAT 3’<br>*mir-51*<br>5’<br>CTTTACATTTTGTTTTCAGAGTCCGAAAAGTCCGTCTACC 3’ | This study | IDT |

| | | |
|---|---|---|
| 5' CAGTTGGAATTCTACGAATGAACTGTATTGCTGCTGGGC 3'<br>*mir-58*<br>5' CTTTACATTTTGTTTTCAGAGCTCGTCATATCCATTGCCC 3'<br>5' CAGTTGGAATTCTACGAATGAGAACAAGTTCGCGAGAGTT 3' | | |
| *mir-34* molecular beacon<br>5' ATCGCGCAACCAGCTAACCACACTGCCTCGCGAT 3' | This study | Sigma Aldrich |
| DCR-1(T961D) repair template:<br>5' CGTCTTAACCTTCTTCAACCTCGAATTCAAAATCAACCACGcCGcTCgAGggaTGTcAGcAACTCCTCAACGTCAAATATTCCTCAAGCATCTGCTTCT 3' | This study | IDT |
| RBPL-1(S367E) repair template:<br>5' TACTTTAGTACAACAGCAAACGACGTTAACATCAGTCAGTgaAGGcACcTCcCTgAGcGCcCAGgtaattagaatttccgtagaatttcaatctgaacttaa 3' | This study | IDT |
| RBPL-1::HA repair template<br>5' tgtttttgaagaattttaatcttagatatttcagAAATCTTCcTCgCGcAAgGTcCCgAAaTACCCATACGATGTTCCAGATTACGCTGAGAGTGTAGATGTCAAACACAAGAGTACCAAGATTAAAT 3' | This study | IDT |
| RDE-4::HA repair template<br>5' AAGAGGCTAAACAGTGTGCTTGTAAATCGGCGATTATCCATTTtAAtACgTAcGAcTTtACcGAcTACCCATACGATGTTCCAGATTACGCTTaAaaatattattgcgtattcctgaaaaatgaagcgtctgaa 3' | This study | IDT |
| DCR-1::FLAG repair template<br>5' CAAATTGAGCAGCAAAGAAGACAAAGCCCATCATTAACAACTGTTGACTACAAGGACGACGATGACAA | This study | IDT |

| |
|---|
| GTAAattatcttcactttctgtgatatgctaagtattaagctatgtgtttctag g 3’ |

**Recombinant DNA / source / identifier**

| | | |
|---|---|---|
| pDD162 (Peft-3::Cas9 + Empty sgRNA) | Bob Goldstein | Addgene plasmid # 47549 ; http://n2t.net/addgene:47549 ; RRID:Addgene_47549 |
| pPD95_77 | Andrew Fire | Addgene plasmid # 1495 ; http://n2t.net/addgene:1495 ; RRID:Addgene_1495 |
| pPD129.36 | Andrew Fire | Addgene plasmid # 1654 ; http://n2t.net/addgene:1654 ; RRID:Addgene_1654 |

**Software and algorithms / source / identifier**

| | | |
|---|---|---|
| NIH ImageJ | Schindelin et al., 2012 | N/A |
| ZEISS Zen 3.7 | ZEISS | N/A |
| MicroChemi | DNR Bio Imaging Systems | N/A |
| GelCapture (Version 7.0.18) | DNR Bio Imaging Systems | N/A |
| TargetScanWorm | Lewis et al., 2005<br>Jan et al., 2011 | N/A |
| ScanSite | Obenauer et al., 2003 | N/A |

| | | |
|---|---|---|
| CFX Manager 3.0 Software | BioRad | N/A |
